# Characterising alphacoronavirus phenotypic traits through diversity-driven selection of spike

**DOI:** 10.1101/2024.11.22.624870

**Authors:** Giulia Gallo, Antonello Di Nardo, Aghnia K. Dewantari, Stephen C. Graham, Dalan Bailey

## Abstract

The capacity for viruses to spillover from one host to another is dependent on their ability to bind to and enter cells from a new host. Using a computational approach that maximises phylogenetic diversity, we selected an optimal subset of 40 alphacoronavirus spike proteins, including the two human viruses NL63 and 229E, and characterised their host-range using broad mammalian APN and ACE2 receptor libraries. Based on this data, we were able to determine molecular genotypes that contribute to receptor tropism and identify alphacoronaviruses with broad (generalist) or restricted (specialist) receptor usage. Strikingly, we observed that generalism and specialism can vary significantly between closely related viruses. Using structural information, we identified key residues that determine the bat tropism of 229E-like viruses, as well as residues in the ACE2 receptor that likely restrict NL63 infection of certain mammals. Furthermore, we observed that the host range of certain bat alphacoronaviruses is expanded by TMPRSS2 priming of spike. All APN- and ACE2-using alphaCoVs in our study interacted with the receptor of at least one animal species within ecological proximity to humans, suggesting new routes for spillover. However, most bat alphacoronaviruses did not use any of receptors in our screen, refining our understanding of coronavirus entry. We propose a new approach to investigating receptor usage for viral orders/families/genera, which is not constrained by focused analysis of a limited number of sequences from human-tropic viruses. Understanding phenotypic traits, such as entry, at this genus-wide level can revolutionise our ability to predict zoonotic potential.

## Introduction

Following the Covid-19 pandemic, there has been a renewed focus on the characterisation and prediction of viruses with zoonotic potential. Between 60% and 75% of human pathogens are believed to have a zoonotic origin^1^ and there is emerging evidence for significant anthropogenic spillover from humans into animals^2^. However, unbiased information on the extant diversity of emerging viruses is scarce, making prediction of spillover potential extremely challenging. For example, the assignment of specific phenotypic traits such as ‘breadth of host receptor usage’ (referred to here as *generalism* and *specialism* for broad and narrow usage, respectively) to genetic level classifications, i.e. species, genus, family, would improve prediction. Unfortunately, data are usually available for only a few species of human viruses, or limited to the study of viral receptors found in humans. Indeed, focused *zoonotic-* rather than broader *spillover-*based research actually restricts One Health impacts, and belies the fact that the next pandemic is likely to be a ‘known unknown’, as SARS-CoV-2 was to virologists in 2019.

The first barrier for any cross-species viral jump is entry into cells of an atypical host, a process reliant on binding of viral attachment proteins to cellular receptors; an interaction which we have characterised previously for single viruses, such as PPRV or SARS-CoV-2. To attempt wider characterisation at the whole genus level, we chose the alphacoronaviruses (alphaCoVs) as a model due to the lack of detailed information on their host-range. For all coronaviruses the spike protein (S) is the major determinant of viral entry, with cleavage by cellular proteases yielding two domains, S1 and S2, that are involved in binding to the cellular receptor and fusion of the viral and cellular membranes, respectively. Additionally, the S2 sub-domain can be further activated at the S2’ site by type II transmembrane serine proteases, facilitating release of the fusion peptide. For human pathogenic betacoronaviruses, proteolytic priming of S2 is associated with increased efficiency of infection in vitro, and augmented severity of pathogenicity in animal models^3^.

To-date, two cellular receptors have been characterized for alphaCoVs: aminopeptidase N (APN) and angiotensin-converting enzyme 2 (ACE2). Human, porcine, canine and feline APN provide entry to alphacoronavirus group I viruses, which includes human CoV 229E^4, 5^, transmissible gastroenteritis virus (TGEV) and the related porcine respiratory coronavirus^6^ (PRCV), as well as some canine and feline coronaviruses^7^. Among alphaCoVs, only the human alphaCoV NL63 is known to use ACE2 as a receptor^8^. Efforts to understand the origin of alphaCoVs, including group I-related viruses, has identified a much broader diversity of virus genotypes in reservoir species such as rodents and bats^9, 10^. Sequencing data from Kenyan bats identified CoVs related to the human alphaCoVs 229E and NL63, indicating that human NL63 might be the result of a recombination event within S between NL63-like viruses of *Triapenos* bats and 229E-like viruses found in *Hipposideros* bats^11^. Taken together, these reports highlight the complexity of alphaCoV evolution, receptor-dependent host range, and support the need for a much deeper understanding of the underlying mechanisms that determine the generalism or specialism of this genus.

To broaden our understanding of the molecular determinants that underpin alphaCov host range, we employed an optimal computational solution to select representative S proteins based on the total phylogenetic diversity. Our goal was to scale down the number of S tested, without losing the richness and heterogeneity of diversity within this genus. In parallel, we built libraries of APN and ACE2 receptors, including relevant peridomestic animals that might act as intermediate hosts for important spillover events. We also selected APN and ACE2 proteins from various bat species to better understand: i) how many *Chiroptera* species can support APN/ACE2-dependent entry of alphaCoV, ii) whether alphaCoVs can transmit among bats sharing the same ecological niche, and iii) how selection pressure might influence viral evolution in reservoir hosts^12^. Receptor usage was assessed using standardised and scalable viral pseudotype entry assays, with our results demonstrating that an unbiased approach is a powerful tool to study virus-host interactions at a genus-wide level.

## Results

### A greedy algorithm allows unbiased selection of alphaCoV Spike representatives

To select S representatives that accurately represent the known diversity of the alphacoronavirus (alphaCoV) genus, we used a greedy algorithm on all S protein sequences retrieved from the ViPR database and previously deposited in GenBank (as of May 2021). Following the downstream analysis pipeline described in Materials and Methods, we selected 40 full-length ORFs from the ∼2200 S sequences initially available for commercial gene synthesis and cloning into mammalian expression vectors. The final number of S sequences was chosen based on our predicted capacity for pseudotyping and receptor screening (Fig. 1a, Table S1). Among these sequences, which included representatives from taxonomically classified sub-genera such as the *Colacovirus*, *Pedacovirus*, *Minacovirus*, *Tegacovirus*, *Nyctacovirus* and *Luchacovirus*, as well as 17 unclassified viruses, the overall level of amino acid conservation was low, especially within the predicted RBD (Fig. 1b, Fig S1). For the most divergent alphaCoVs, two unclassified viruses and viruses classified within the *Soracivirus* and *Luchacovirus* sub-genera, an RBD could not be readily identified. The majority of the selected S proteins are from viruses isolated in bats (27/40); however, better characterised and/or more well-established representatives of alphaCoVs were also greedy-selected, including those that infect domesticated animals, e.g. CCoV and PEDV, along with two endemic human alphaCoVs, hCoV/229E and hCoV/NL63. To confirm that cloned, codon-optimised S ORFs could efficiently produce S proteins that were pseudotyped onto HIV-1 based lentiviruses, pseudoviruses were purified and S incorporation confirmed by immunoblot (Fig. S2 and summarised in Fig. 1b; ‘PV?’ column). For the six S proteins that could not pseudotype, we attempted to substitute them for S proteins that have been reported to pseudotype and that share at least 95% amino acid similarity (Fig. S3). However, the only match was for PEDV/KDJ, which was replaced by PEDV/Colorado^13^. The selected human 229E S protein sequence pseudotyped to low levels and was not functional in our downstream applications (Fig. S5), therefore we replaced it with a different strain of the same species.

**Figure 1.**
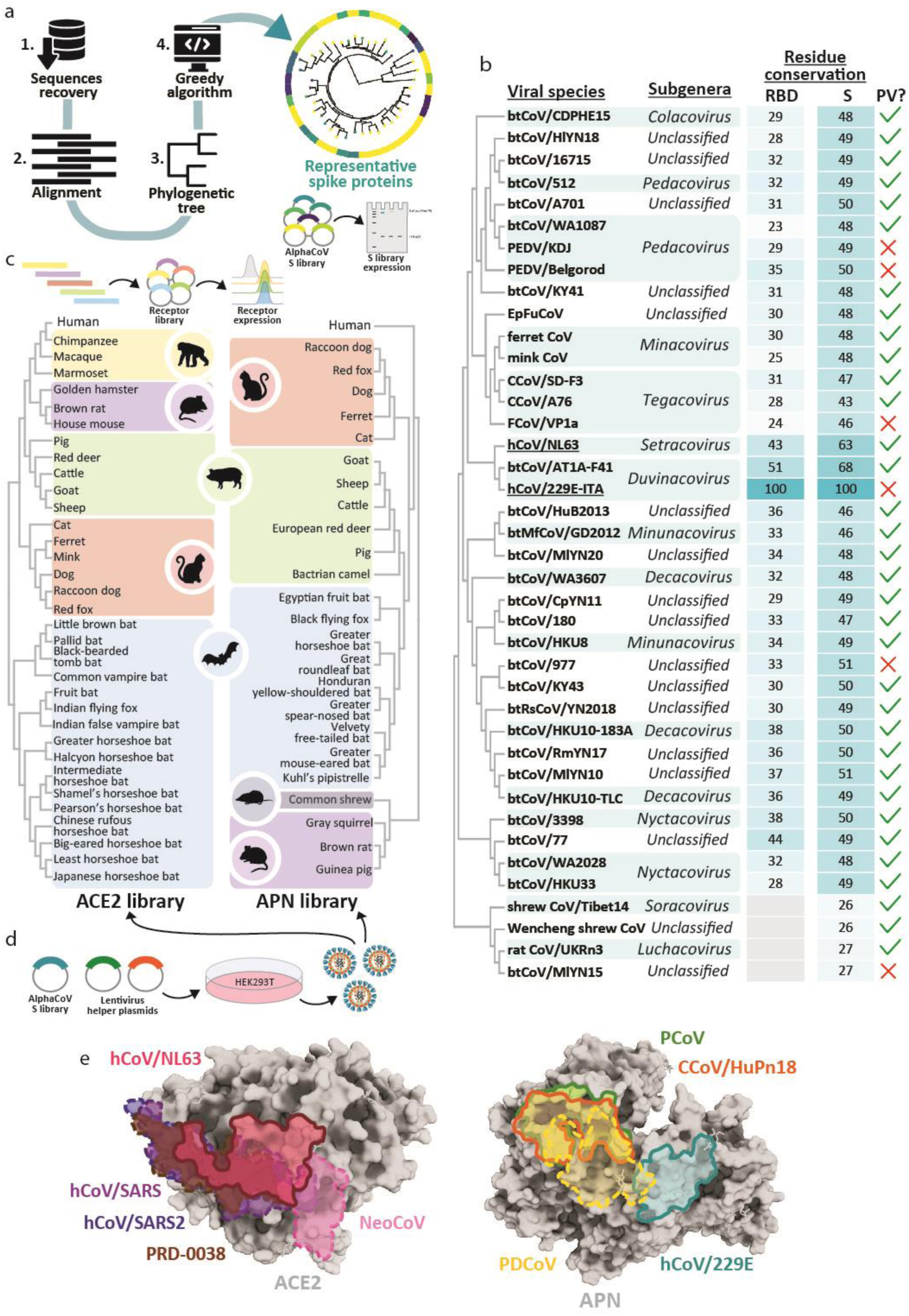
Spike and receptor library information and RBD-receptor binding interfaces. (a) Scheme summarising the selection process used to create the Spike library. Full-length amino acid sequences of alphaCoVs were recovered from public repositories (1.) and aligned (2.) to generate a phylogenetic tree (3.), which was used to estimate the sum of branch lengths distance between all tips, in a pairwise manner, and select 40 representatives Spike (S) sequences using an optimal greedy-based algorithm (4.). The suitability of these proteins to be incorporated in lentiviral pseudotypes has been assessed by immunoblot. (b) General information about the sequences, such as their ICTV subgenus classification and their percentage of identity of the RBD and the full-length S, compared to hCoV/229E. Underlined names denote viruses that are endemic in humans. Additionally, successful pseudovirus (PV) incorporation is reported. (c) The expression of the APN and ACE2 libraries was assessed by flow cytometry. Animal species included in the libraries are listed, categorized by their mammalian order. (d) Scheme of procedure for alphaCoV host range screening, which relies on alphaCoV PV production using lentiviral helper plasmids. (e) Footprints of experimentally validated coronavirus RBDs on their receptor are illustrated on human ACE2 and canine APN. Sarbecoviruses hCoV/SARS, hCoV/SARS2 and clade 3 bat CoV PRD-0038 RBDs bind similar ACE2 surface residues (PDB: 2AJF^41^, 6M0J^42^ and 8U0T^43^, respectively). However, the RBD of MERS-related NeoCoV, identified in bats, interacts with a different region of ACE2 (PDB: 7WPO^44^). Binding of hCoV/NL63, the only alphaCoV known to use ACE2 (PDB: 3KBH^45^), partially overlaps the binding regions of sarbecoviruses. hCoV/229E binds to a distinct region of APN, compared to PCoV/PRCV and CCoV/HuPn18 (PDB: 6ATK^46^, 4F5C^6^ and 7U0L^47^, respectively). DeltaCoV PDCoV footprint on APN (PDB: 7VPQ^48^) partially overlaps hCoV and CCoV/PCoV binding sites. AlphaCoV binding footprints are in full lines, while betaCoV and deltaCoV footprints are drawn with dashed lines.

To date, APN and ACE2 are the only known alphaCoV receptors involved in entry. Concurrent to the establishment and validation of this S library, we also developed an equivalent plasmid-based APN and ACE2 expression library, representing 25 and 34 mammalian species, respectively (Fig. 1c, Tables S2-S3). Of note, greedy-selection was also used to return animal species for receptor libraries; however, selected sequences were not ecologically and/or biologically relevant based on the current knowledge of alphaCoVs, therefore we selected the representative species manually. Where possible, we included the species wherein individual viruses were identified, as well as domesticated and peridomestic animals (which might bridge the gap between unknown reservoirs and humans). We also selected genetically divergent bat species, to account for the reservoir biology of coronaviruses. We verified the expression of these tagged receptors libraries by flow cytometry (Fig S4) and proceeded with pseudovirus infection of cells transiently expressing individual receptors from our libraries (schematically depicted in Fig 1d). Coronavirus binding to ACE2 has been extensively characterized, showing that most RBDs bind a generally conserved surface domain, except for bat NeoCoV (Fig 1e). For example, the binding surface of hCoV/NL63, the only established ACE-using alphaCoV, partially overlaps those of SARS, SARS-CoV-2 and bat PRD-0038. Elsewhere, it has previously been shown that 229E interacts with a different region of APN compared to canine (CCoV) and porcine (PCoV) alphaCoVs, which bind overlapping interfaces. The footprint of porcine deltacoronavirus (PDCoV), which also uses APN for entry, partially superposes on both the hCoV/229E and CCoV/PCoV interaction interfaces (Fig 1e).

### APN and ACE2 receptor usage is relatively rare amongst alphaCoVs, and these viruses exhibit differing degrees of host-range generalism and specialism

To facilitate receptor identification, we assayed whether the pseudotyped S from our alphaCoV library could use any APN and/or ACE2 from our receptor libraries (Fig. 2a and Fig. S5-S9). Building on our understanding of protease activation of S2’ during entry, all experiments were performed in the presence or absence of human TMPRSS2. For brevity, the total number of APNs or ACE2s used by each S for entry is summarised in Fig. 2b, as an overall indicator of their receptor usage traits (generalist vs specialist). As expected, hCoV/229E and CCoVs could enter using APN, while hCoV/NL63 used ACE2. Despite a feline CoV being included in the library, it did not pseudotype, and we could not find any report identifying a FCoV S to use as replacement for entry assays. For PEDV, a substitute was identified, but it did not use porcine APN in our experimental settings. To provide confidence that our porcine APN was functional, we pseudotyped another porcine alphaCoV, PRCV, and showed it did use this cognate receptor for entry (Fig. S10). Surprisingly, the majority of the other alphaCoVs could not use any of the expressed APNs or ACE2s to enter cells, regardless of co-expression with TMPRSS2 (Fig. S5-S8). Nevertheless, we did find two bat-origin alphaCoV, WA1087 and AT1A-F41, that used APNs to enter cells, albeit not human APN. To our knowledge this is the first study reporting APN-usage by alphaCoVs isolated in bats (Fig. 2). Based on the number of receptors used and the genetic relationship of the host APNs or ACE2s, alphaCoV were divided into four categories: *multi-order specialists*, such as hCoV/229E and btCoV/WA1087, which can use the APNs from a small number of species from distinct animal orders; *true specialists*, like CCoVs, that can only enter cells expressing the APN from species within only one mammalian order; *broad generalists*, e.g. hCoV/NL63, which interacts with the ACE2 from multiple species within various orders; and *order generalists*, such as btCoV/AT1A-F41, which interacts with receptors from many animals within the same mammalian order (Fig. 2b). Of note, we also examined if any S in our alphaCoV library could use human TMPRSS2 or human DPP4 alone to enter cells, since they act as receptors for the betacoronaviruses HKU1 and MERS (respectively), but we could find no evidence for entry (Fig. S11). These findings suggest that: (i) APN or ACE2 usage is a relatively rare phenomenon within the alphaCoV genus; and (ii) the spectrum of host-range receptor usage (generalism vs. specialism) is highly variable within the smaller sub-group of APN- or ACE2-using coronaviruses.

**Figure 2.**
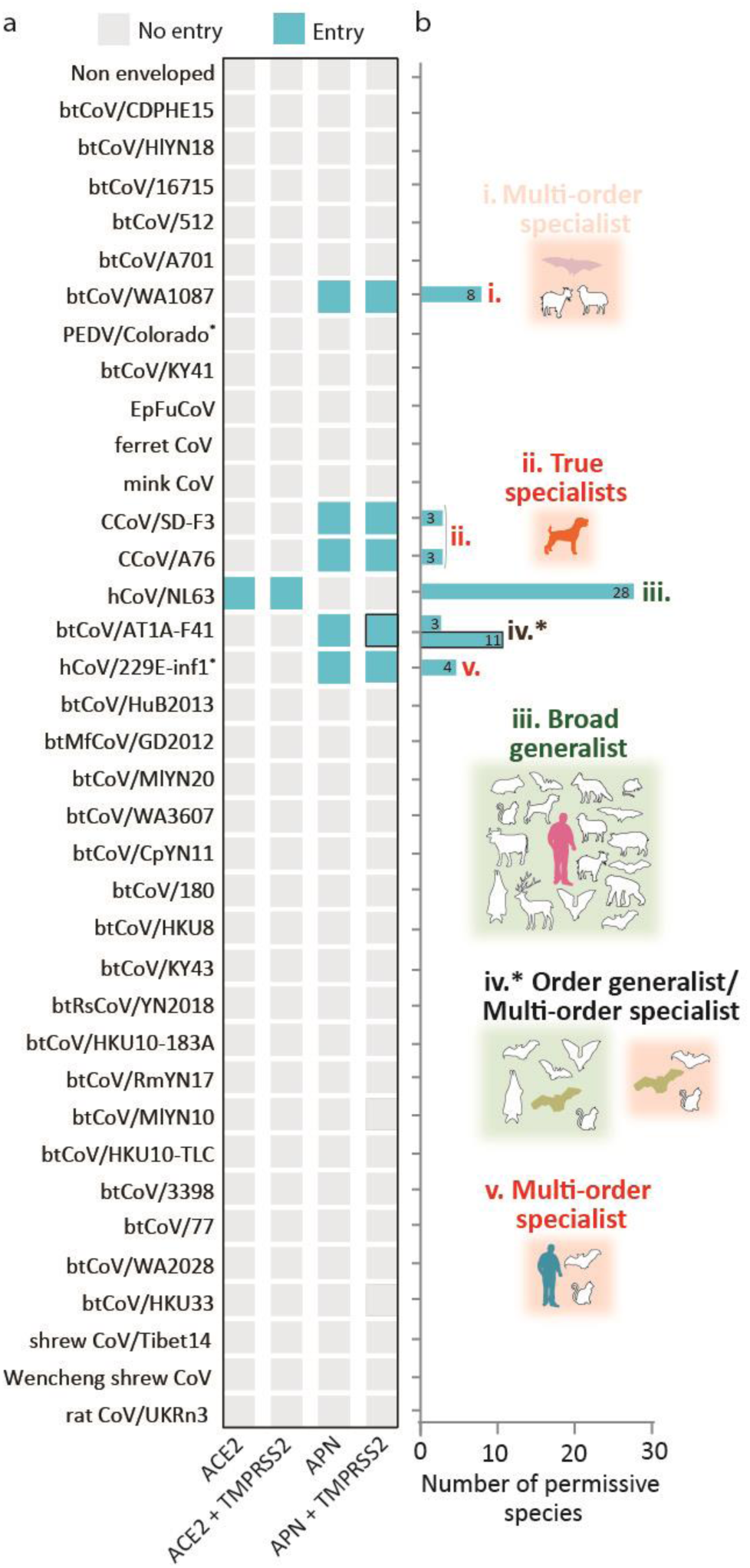
AlphaCoV S usage of ACE2 and APN libraries, in absence or presence of cellular transmembrane protease TMPRSS2. (a) Screening of receptors using pseudotyped alphaCoV S. Cyan indicates that the receptor of at least one species is permissive for entry, while grey denotes no entry. Sequences that did not pseudotype and were replaced by others described as functional in the literature are marked using a star symbol (*). Positive results were reproduced at least three times, using independent biological replicates. The average of these results is shown in Figures S5–S8. All experiments were carried out in technical triplicates. (b) For the positive results, the number of animal species whose APN or ACE2 permitted entry is shown. Based on the intra- and inter-order number of permissive animal species, alphaCoVs were classified in four categories: true specialists (CCoVs), which use the receptor of only few species belonging to the same mammalian order; multi-order specialists (btCoV/WA1087 and hCoV/229E-inf1), that use the APN of a small number of species of different animal orders; order generalist (btCoV/AT1A-F41), which use a wide selection of species of the same order; and broad generalist (hCoV/NL63), that interact with a plethora of ACE2 receptors from different mammalian orders.

### Human alphaCoVs NL63 and 229E bind to different receptors and exhibit differing host range traits

Pseudotype analysis identified hCoV/NL63 S as a broad generalist with regards to its ACE2 usage (27/34 permissive species) (Fig. 3a). To understand this generalism in more detail we aligned the ACE2 sequences from non-permissive species and compared them to the consensus found in ACE2s from permissive species. Residue conservation across our ACE2 library was projected onto the structure of ACE2, and the binding site of hCoV/NL63 RBD highlighted (Fig. 3b; pink outline, right panel), identifying that residue 354 is poorly conserved within this domain (Fig. 3b). At this position, a conserved glycine found in permissive species is replaced by extended polar residues in the ACE2s of non-permissive marmoset (Q; New World monkey), ferret and mink (R/H; *Mustelidae*), and *Coelophyllus* horseshoe bats (D/N; Shamel’s and Pearson’s horseshoe bats) (Fig. 3b). ACE2 position 354 is involved in a network of interactions with the hCoV/NL63 RBD, including RBD residues Y498, Y499, W585 and G537 (Fig. 3c). Interestingly, G354 is also at the interface of interaction with SARS-CoV-2 RBD. However, it does not seem to explain the host range based on binding assay of mammalian ACE2s and RBD. While ACE2 residue 354 is at the core of the interaction with hCoV/NL63, this residue is at the periphery of the binding interface with SARS-CoV-2 RBD (Fig. S12). To examine this restriction further we substituted R354 and Q354 to glycine in ferret and marmoset ACE2, respectively, and showed that these mutants supported efficient entry of hCoV/NL63 pseudotypes at a level similar to human ACE2 (Fig. 3c). An equivalent analysis was undertaken for the APN using alphaCoV hCoV/229E. Only human, cat and velvety free-tailed bat APN were permissive to pseudoparticle entry by the multi-order specialist hCoV/229E (Fig. 3d), with camel being significantly more permissive in the presence of TMPRSS2 (3% and 73% -/+ hTMPRSS2, discussed below). Since many species APNs cannot confer entry to cells, identifying specific residues that could account for host range is more challenging. However, upon similar alignment of the APN sequences from our library and the projection of conservation onto the surface of an APN structure, highlighting the established hCoV/229E RBD binding site. One of the most variable motifs is contact residues D288 and Y289 in human APN (Fig. 3e), which interact with hCoV/229E RBD residues C317, G314 and K316 (Fig. 3f). We confirmed the importance of this interaction by mutating D288K and Y289S of human APN to account for the variability observed in the non-permissive carnivore (D288K) and ruminant (Y289S) species (Fig. 2d and 2e). Consistent with our sequence analysis, both these substitutions significantly affected the entry of 229E pseudotypes.

**Figure 3.**
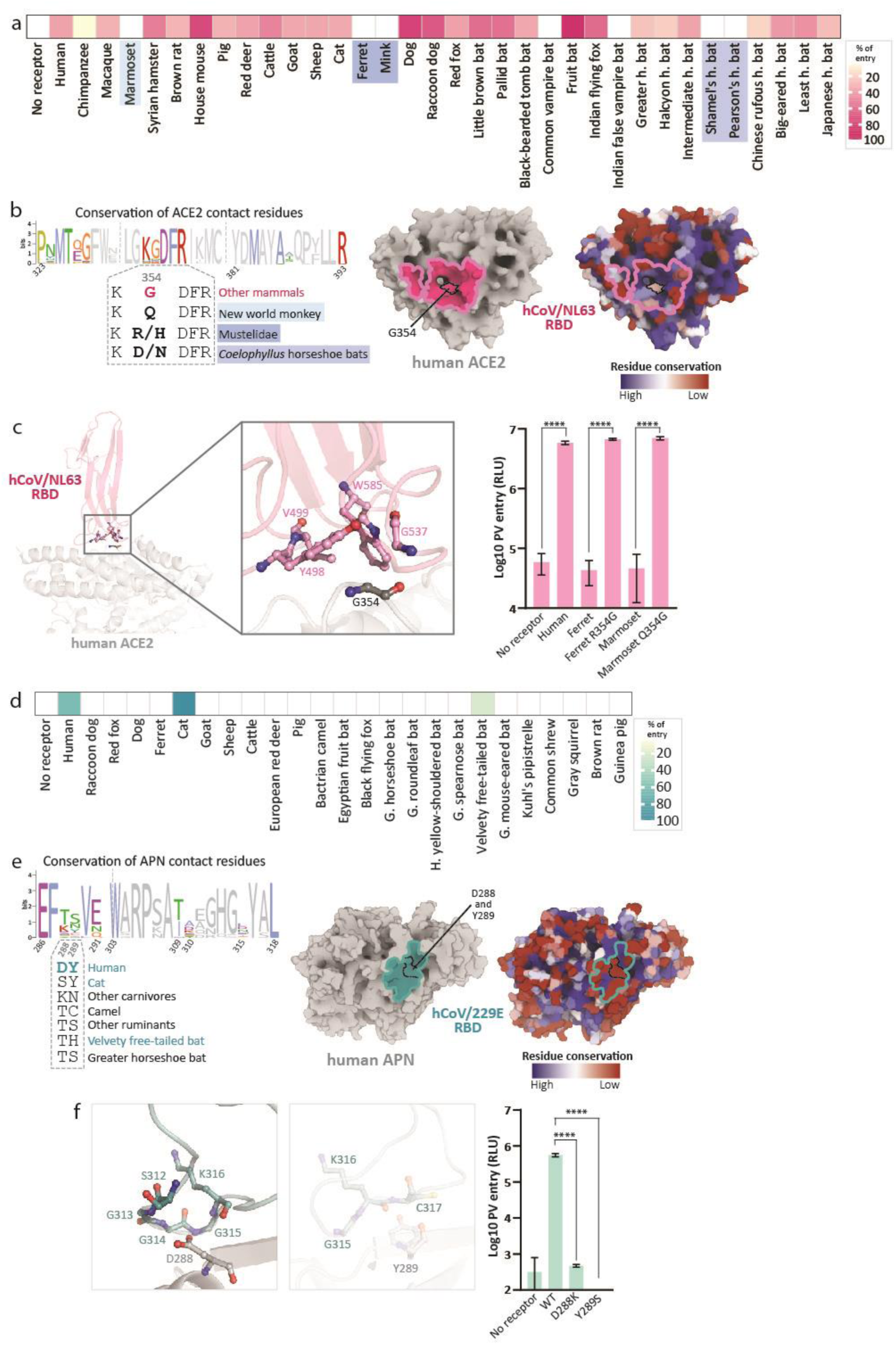
Animal host range of human alphaCoVs NL63 and 229E with their respective receptor, ACE2 and APN, and their molecular determinants of interaction. (a) Pseudotyped NL63 S was used to assess usage of ACE2 of different mammalian species. Raw data is normalised using min-max feature scaling and shown as percentage of usage. Almost all ACE2 tested could support entry by NL63. An average of three biological replicates is shown. Restricted species are highlighted in shades of blue. (b) The footprint of NL63 interaction with ACE2 (PDB: 3KBH45) is highlighted in magenta (middle) and ACE2 residue conservation in the receptor library is illustrated (right). A WebLogo of ACE2 sequences, with coloured residues representing those at the interface with NL63 RBD, is shown (left). To compare the sequence of restricted versus not-restricted species, an alignment of residues 353-357 is shown. Correlation between the entry assay and the alignment highlights residues that are responsible for host range, such as G354. (c) NL63 RBD loops that interact with ACE2 (PDB: 3KBH45), with the residues in proximity of ACE2 G354 shown. Following site-directed mutagenesis to substitute residue 354 of ferret and marmoset, the contribution of G354 to NL63 permissivity was assessed by entry assay. Substitution to glycine of the equivalent residues in ferret and marmoset ACE2s conferred entry, providing a molecular explanation for restriction. (d) Pseudotyped 229E-inf1 was used to test entry with the APN receptor library. In contrast to the broad host range of NL63, the APN proteins of few species support entry of 229E. Raw data is rescaled as percentage of usage. Experiments were repeated with three biological replicates, and an average of those is shown. (e) Footprint of 229E on human APN (PDB: 6ATK46) is coloured in teal (middle) and APN residues conservation in the receptor library is illustrated (right). A WebLogo of APN residues 286-318 shows per-residue sequence conservation (left). Residues in contact with 229E RBD are coloured. Variability of residues 288-289 of different species is shown. (f) The 229E RBD residues in contact with hAPN D288 and Y289 are shown on the RBD-receptor structure (PDB; 6ATK46). Pseudoviruses were used to assess the contribution of D288 and Y289 to 229E entry by site-directed mutagenesis, showing them to be essential for 229E entry. For statistical analysis, values of entry with mutated receptors were compared to the corresponding WT using ordinary one-way ANOVA. Experiments were repeated at least two times, in technical triplicate.

### The host ranges of 229E and 229E-like alphaCoVs are determined by single residues within the S RBD

In addition to hCoV/229E, another *Duvinacovirus* was greedy-selected - the 229E-related virus btCoV/AT1A-F41, isolated and sequenced from Aba roundleaf bat (*Hipposideros abae*) in Ghana and Cameroon^14, 15^. Although the btCoV/AT1A-F41 S and RBD share only 68% and 51% sequence identity relative to hCoV/229E, respectively, we showed that both viruses utilise APN to allow entry and both are multi-order specialists. However, their APN host-ranges are different: btCoV/AT1A-F41 does not use human or camel APN but can utilise a wider range of bat APN receptors (Fig. 4a). We compared the S and RBD loops from additional 229E-like viruses derived from human, camel and bats, and noticed that the N-terminal domain (NTD) varies considerably in length for these viruses (Fig. 3b). This region is upstream of the RBD, which is found in the C-terminal region of S1. It is postulated that the loss of the NTD in human 229E could have been involved in previous host and organ tropism switches^16^. To examine this further, we deleted the NTD of btCoV/AT1A-F41, to assess its contribution to host range.

**Figure 4.**
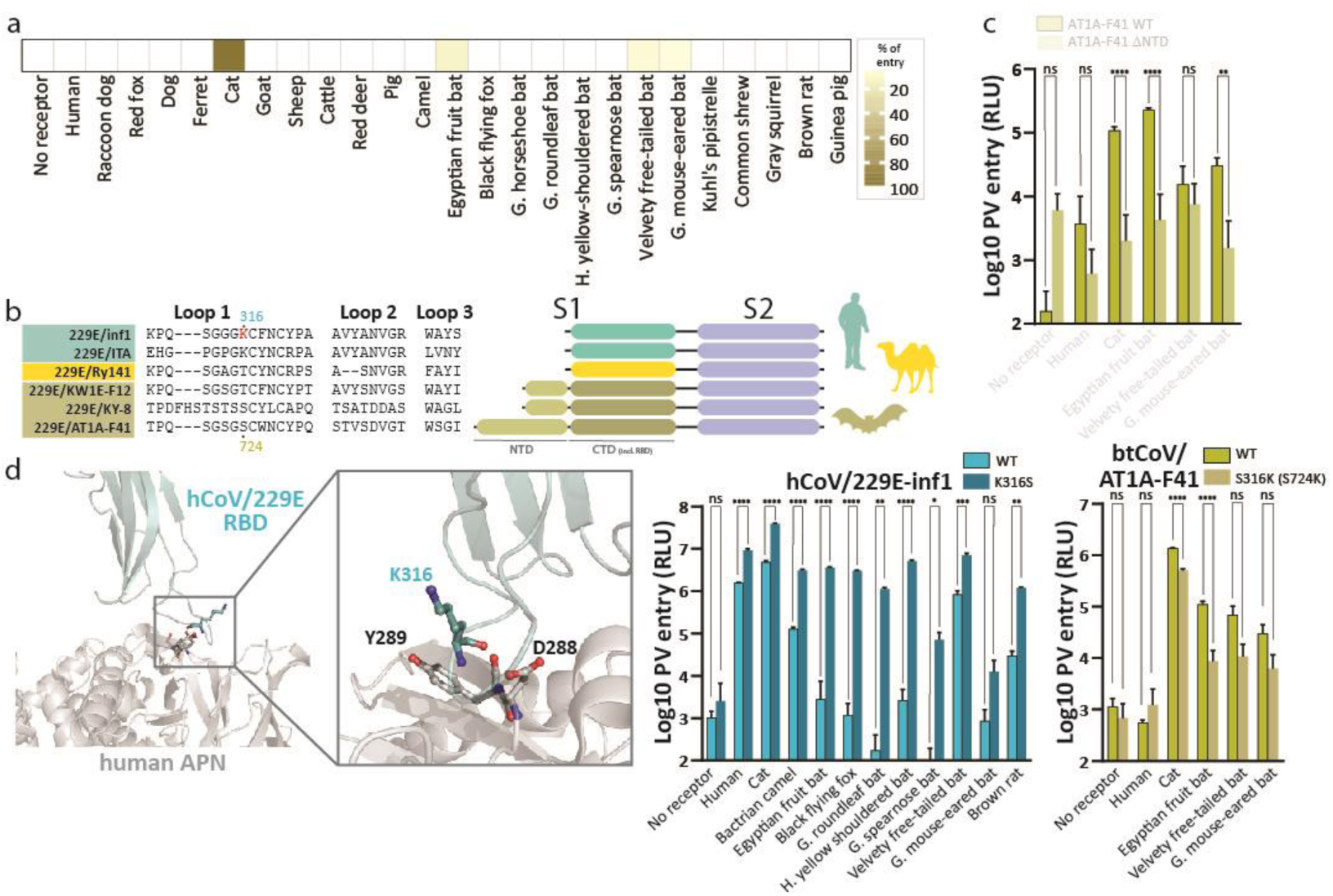
Animal host range of 229E-like bat alphaCoV AT1A-F41, and the impact of RBD substitutions on 229E viruses on host range. (a) Pseudotyped btCoV/AT1A-F41 was used to assess its host range based on usage of APN receptor. The APN host range is limited, similarly to what has been observed with human 229E. (b) Alignment of RBD loops of different 229E-like viral species found in human (teal), camel (yellow) and bat (brown) hosts (Genbank ID of the camel and bat S sequences not included in the S library: 229E/Ry141, YP_009194639.1, 229E/KW1E-F12, ALK28771.1, 229E/KY-8, APD51507). On the right, their S gene organization is showed, highlighting the absence of a N-terminal domain (NTD) in human and camel 229E sequences, and the longer NTD of bat AT1A-F41. (c) To assess the contribution of AT1A-F41 NTD to its entry using APN, the domain was removed and the S pseudotyped. The protein lost the ability to provide entry. Statistical analysis was performed using two-way ANOVA with Šidák multiple comparison test. (d) Structure of 229E RBD-hAPN, where the residue of hAPN in proximity to 229E-RBD K316, D288 and Y289, are shown (PDB: 6ATK^46^). Based on RBD alignment, the K316S substitution was introduced in human 229E, showing an increase in the number of APN species that support entry by human 229E. The complementary substitution was inserted in AT1A-F41 S, and the ability of pseudovirions to enter cells was lost. Statistical analyses were performed using multiple unpaired t test, comparing WT to mutated S.

However, we found that the btCoV/AT1A-F41S lacking the NTD was non-functional, since previously permissive APNs, such as cat, Egyptian fruit bat and great mouse-eared bat could no longer confer entry, and there was no corresponding increase in human APN usage (Fig. 4c). More focused analysis on the loop 1 alignment identified that the aforementioned RBD position K316 in hCoV/229E is conserved only in human-tropic viruses, whereas it is substituted to short polar residues in bat 229E-like viruses (serine/threonine). As discussed earlier, this residue is in close proximity to residues D288 and Y289 in hAPN that are critical for utilisation of APN by S (Fig. 3f). To understand the role of this residue in host-range, we mutated hCoV/229E and btCoV/AT1A-F41 S, introducing K316S and S724K (the corresponding residue in this NTD-containing alphaCoV S), respectively, and analysed entry using our APN library (Fig. 4d, Fig. S13). Interestingly, the K316S mutation in hCoV/229E extended the host range of this S protein, significantly widening the usage of a range of bat APNs, without compromising human APN tropism. On the other hand, the equivalent S724K substitution in btCoV/AT1A-F41 reduces the host range, with only cat APN entry being maintained. Furthermore, it highlights that single amino acid mutations in APN-using alphaCoVs, such as 229E, can extend the host-range traits of this relative specialist to a more generalist phenotype; highlighting a potential risk for reverse zoonosis/anthropogenic spillover.

### The host range of certain bat-derived alphaCoVs is expanded by TMPRSS2 priming of the S protein

It is well established that priming of coronavirus S proteins at the S2’ site by the serine protease TMPRSS2 enhances the efficiency of entry in cells expressing APN/ACE2/DPP4^17^. Building on our earlier observation that the range of APNs used by btCoV/AT1A-F41 increased with TMPRSS2 (Fig. 2b), we examined this result in a broader context through comparison with other alphaCoVs that can use APN. In addition to btCoV/AT1A-F41, we identified another bat alphaCoV (WA1087) that could interact with APNs to enter cells, although it too had a receptor usage profile distinct from 229E (Fig. 2 and Fig. 5a). We better dissect this by further comparing the APN host range of btCoV/WA1087 and btCoV/AT1A-F41 with hCoV/229E in the presence or absence of human TMPRSS2 (Fig. 5a). We found that the host range and entry efficiency of btCoV/WA1087 was not impacted by the co-expression of hTMPRSS2. In contrast, 229E-related S proteins showed significant differences: in the case of hCoV/229E, priming of S2’ altered the relative efficiency of entry, with human APN becoming the receptor associated with the highest entry signals, compared to cat in the absence of hTMPRSS2. Surprisingly, for btCoV/AT1A-F41, the priming of S broadened the host range, with numerous additional bat APNs able to confer entry to cells. In addition, the primed S now used velvety free-tailed bat APN as its most efficient receptor, compared to cat in the absence of hTMPRSS2 (Fig. 5a). Focusing on relevant hosts, we again confirmed that the S priming of btCoV/WA1087 did not confer appreciable enhancement of entry to this pseudotype, whereas hTMPRSS2 increased entry of hCoV/229E and btCoV/AT1A-F41 by 10-fold (Fig. 5b). Alignment of the putative S2’ regions targeted by serine proteases showed that btCoV/WA1087 does not possess an arginine residue corresponding to the hCoV/229E cleavage site R685, but btCoV/AT1A-F41 does (Fig. 5c). To confirm that the phenotypic differences in entry observed upon co-expression of TMPRSS2 were due to this enzyme’s catalytic activity, and not its role as a potential attachment co-factor, we repeated these assays in the presence of a catalytically inactive hTMPRSS2 mutant, S441A^18^. These data demonstrated a catalytically active hTMPRSS2 is required to increase and/or widen the entry phenotype of 229E-related viruses (Fig. 5d, Fig. S14). Interestingly, one consistent exception was cat APN, whose usage by alphaCoVs appears less sensitive to co-expression of TMPRSS2. Even though btCoV/WA1087 seemed to enter with greater efficiency in the presence of the inactive TMPRSS2, only results with >10-fold change were considered as being significant for these entry assays. Finally, we examined the activity of another serine protease, human HAT, to assess its ability to extend the host range of the S of btCoV/AT1A-F41. Similar data to hTMPRSS2 were obtained, with hHAT broadening APN usage to additional bat species (Fig. 5e, Fig. S14). These data provide compelling evidence for serine protease-based cleavage of S2’ in extending the host range of alphaCoVs, which could be attributed to the presence of basic residues in the target site.

**Figure 5.**
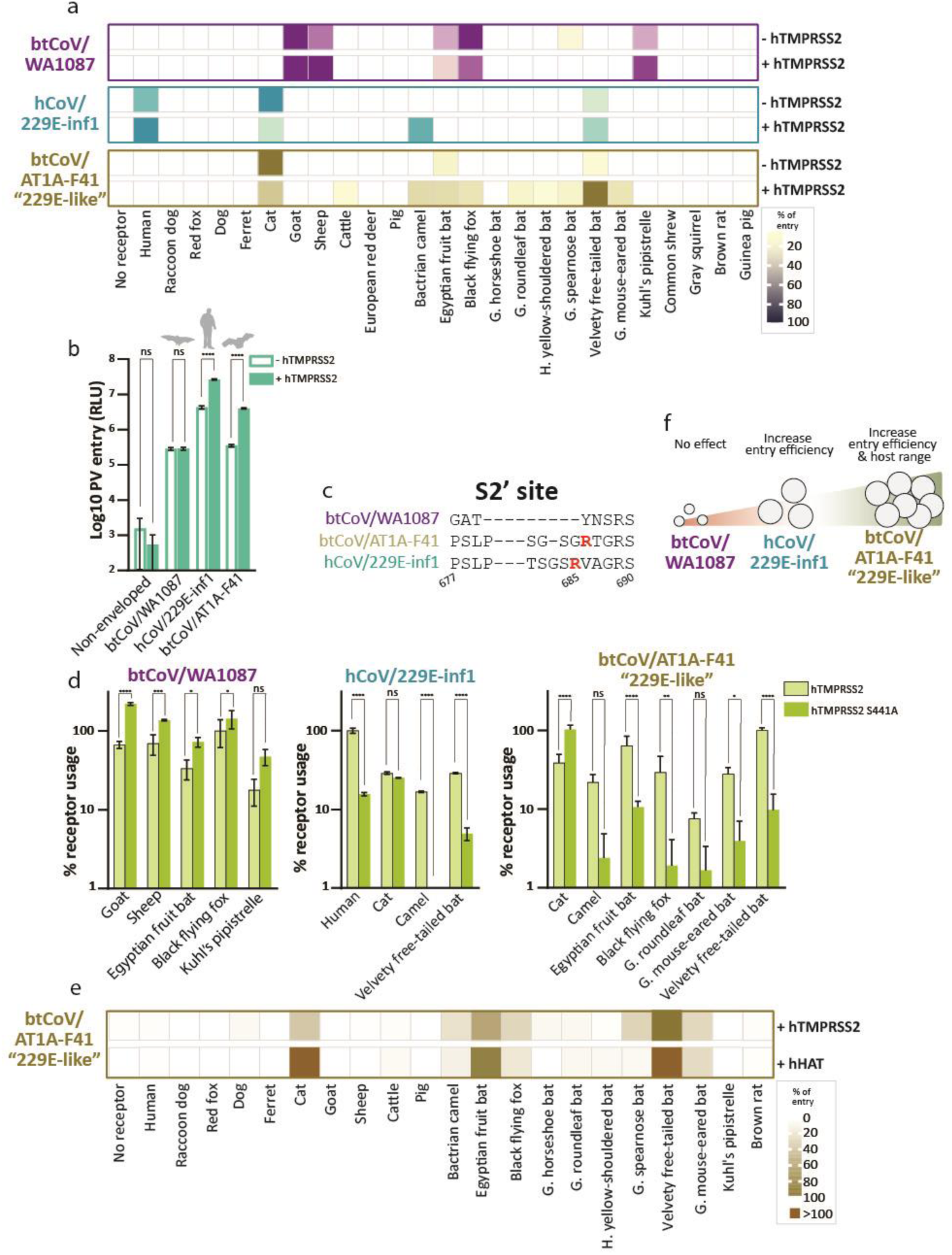
Host range of WA1087, an alphaCoV using APN, and the impact of cellular proteases priming APN-tropic bat-derived alphaCoV S. (a) Effect of human TMPRSS2 on viral host range, assessed by entry assay using pseudotyped S. Co-expression of TMPRSS2 did not affect btCoV/WA1087 host range. However, in the case of 229E-like viruses, TMPRSS2 impacted the efficiency of binding to certain APNs (hCoV/229E-inf1) or increased the number of species that support entry by S (btCoV/AT1A-F41). (b) Entry raw data showing the impact of hTMPRSS2, which was co-expressed with black flying fox APN, human APN or velvety free-tailed bat APN for btCoV/WA1087, hCoV/229E-inf1 and btCoV/AT1A-F41, respectively, as compared to APNs only. Increased efficiency of entry was observed only for 229E-like viruses. Statistical analysis was performed using two-way ANOVA with Šidák multiple comparison test. (c) Alignment of the predicted cleavage site of TMPRSS, showing absence of the basic residues at the S2’ cleavage site of btCoV/WA1087. (d) To discern if TMPRSS2 impact on entry was due to its involvement as a potential co-receptor or for its enzymatic activity, we co-expressed a catalytically inactivated form of human TMPRSS2 along with different APNs. While no different was observed for btCoV/WA1087, the inactivated form reduced entry of hCoV/229E and btCoV/AT1A-F41 by around 90% for all except cat APN. (e) To further validate the increased APN host range of btCoV/AT1A-F41 due to TMPRSS2 activity, we repeated the entry assay with another serine protease, human HAT. A similar increase in entry efficiency and host range was observed. (f) We observed three scenarios when priming bat-derived alphaCoV S proteins with serine proteases: in case of btCoV/WA1087, cleavage at the S2’ site is not possible and TMPRSS2 does not alter entry efficacy; for hCoV/229E, priming of S2’ increases the efficiency of entry; in case of btCoV/AT1A-F41, the presence of serine proteases not only increases entry, but also broadens the APN-dependent host range. Two-way ANOVA analysis with Dunnett’s multiple comparison test was used for statistics.

### The diversity of APN usage by canine alphaCoVs isolates is complex and not easily defined by individual substitutions at the RBD-APN interface

Our optimal approach to alphaCoV S selection returned two canine coronaviruses, CCoV/A76 and CCoV/SD-F3, classified within the *Tegacovirus* sub-genera. Despite sharing only 67% (full S) and 53% (RBD only) amino acid identity, they showed very similar APN usage patterns, namely being true specialists restricted to only the canid APNs (dog, red fox and raccoon dog), regardless of the presence/absence of serine protease (Fig. 2a and 6a). To extend our analysis to reflect the recent epidemiology of CCoV, we added two more tegacovirus S proteins that are considered the etiological agents of hospitalized cases of pneumonia in humans: CCoV/HuPn18^19^ and CCoV/Z19^20^. Interestingly, upon their assessment in equivalent receptor usage screens, we demonstrated that their APN-dependent host range was broader and included multiple animals from various orders but, significantly, not human APN. Similarly to the restricted CCoVs, the host range of these more generalist viruses was not affected by the presence of hTMPRSS2. CCoV/A76 is the most divergent S, while CCoV/SD-F3 shares 78% and 81.5% sequence identity to the generalist CCoVs (full-length S and RBD, respectively). Of note, the RBD of HuPn18 and Z19 is identical (Fig. 6b). To examine a role for RBD sequence variability in determining CCoV host range, we projected the relative amino acid conservation of these isolates on to the structure of the HuPn18 RBD, focusing on the two interacting loops, 1 and 2, that bind directly to APN (Fig. 6c; top panel). Similarly, we constructed a conservation map for our APN library and projected this on to the canine APN structure, highlighting the RBD binding footprint on this receptor (Fig. 6c, bottom panel). We then compared this to an alignment of the RBD sequences of the four CCoVs in our library (Fig. 6d). To assess the overall contribution of the RBD and the interacting loops to host range, we first produced chimeric S between SD-F3 and HuPn18 (Fig. 6e). Initially, we swapped the complete RBD between SD-F3 and HuPn18 S proteins and assessed entry of these two chimeras compared to parental S sequences. We found that, although substituting the specialist SD-F3 RBD into the generalist HuPn18 S backbone reduced the host range, there was not a complete phenotypic switch, since cattle and velvety free-tailed bat APN usage was lost, but camel and cat APN usage was retained. SD-F3 chimeras containing the RBD of HuPn18 were less informative, since the resultant pseudotyped S could not use efficiently use any APN for entry (Fig. 6f). Considering that loop 3 of the specialist SD-F3 and generalists HuPn18 and Z19 are identical, we focused our analysis on loops 1 and 2 of SD-F3 and HuPn18 by building individual loop chimeras to evaluate their role in APN usage and host range. Of note, these loops contribute most of the contact interface between RBD and APN. Even though loop 2 seems to be less conserved than loop 1 among CCoVs (Fig. 6c top), loop 2 chimeras of HuPn and SD-F3 had no effect on the host-range of the backbone (Fig. 6g). Conversely, we observed that loop 1 chimeras were less well tolerated: insertion of the HuPn18 loop 1 into SD-F3 S reduced the overall level of entry conferred by the canid APNs dog and red fox. Similarly, an SD-F3 loop 1 in a HuPn18 S backbone affected the usage of all non-canid APNs, markedly so for velvety free-tailed bat APN. Since modification to loop 1 had the most dramatic effects on APN usage, we prepared single amino acids mutants, based on residue conservation in this motif (Fig. 6d). However, making specialist-specific substitutions R540K and P550L in the generalist HuPn18 had no effect on APN usage or host range. Similarly, the equivalent substitutions in SD-F3 (K547R and L557P) had almost no effect on SD-F3 host range although they did decrease the overall efficiency of entry with the canid APNs we examined (Fig. 6h). These data highly support the hypothesis that receptor usage and host range can vary significantly between viruses classified within the same species, and that the genetic determinants of generalism and specialism are not always easily definable to the RBD-receptor contact interface alone, and perhaps are the result of changes elsewhere in the S protein.

**Figure 6.**
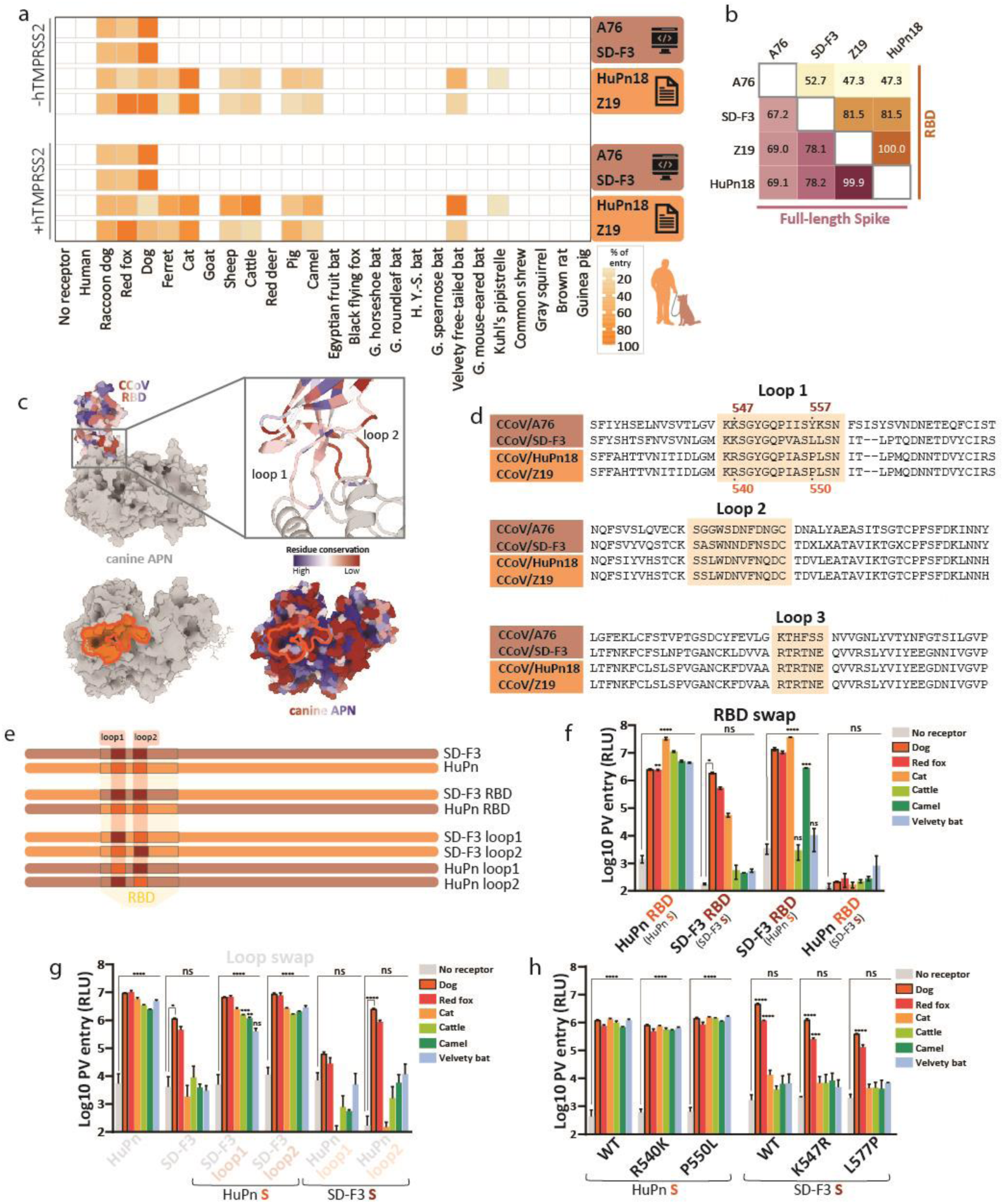
Animal host range of canine alphaCoVs and the contribution of their RBD, or its subdomains, to APN usage. (a) Pseudotyped canine alphaCoVs were used to assess their host range based on usage of APN receptor. Two Spikes, A76 and SD-F3, were greedy-selected, while Z19 (GenBank: QWY12682) and HuPn18 (GenBank: QVL91811) were included due to their reported relevance as zoonotic viruses. The latter show a broader host range, as compared to A76 and SD-D3. Moreover, the presence of human TMPRSS2 did not alter APN usage. (b) Amino acid identity (%) of the canine Spike RBD or full-length protein. Although they were isolated from different part of the world, HuPn 18 and Z19 share almost 100% sequence identity. A76 is the most divergent sequence, especially its RBD. (c) The footprint of HuPn18 RBD binding region on canine APN (PDB: 7U0L^47^) is coloured in orange on its structure (left) and APN residues conservation in the receptor library is illustrated (right). (d) Alignment of canine alphaCoV RBDs, with APN-proximal loops highlighted in orange. (e) Schematics of SD-F3 (dark red) and HuPn18 (orange) chimeras produced. Either the RBD, or the individual loops, were swapped and used for entry assay with pseudotyped chimeric S. (f) Results of entry assay with CCoV swapped RBDs. The introduction of SD-F3 RBD in HuPn18 background (SD-F3 RBD) partially impacted the host range of HuPn18, as cattle and velvety bat APN could no longer support entry. However, camel APN could still provide entry. For the complement construct – HuPn18 RBD in SD-F3 background (HuPn18 RBD) – entry with all receptors was lost, suggesting poor tolerance of S manipulation for canine alphaCoV with restricted host range. (g) Entry data reveals the impact of individual loop substitutions on APN permissivity: while the substitution of SD-F3 loops in HuPn18 S only have a mild effect on the efficiency of entry, the swap of HuPn18 loop1 in SD-F3 drastically reduced entry. The introduction of HuPn loop2 in SD-F3 had no effect on entry. (h) Entry assay testing individual substitutions in the loop 1 of SD-F3 and HuPn18. No effect on host range was observed for either of the viruses. Significance was assessed using two-way ANOVA with Dunnett’s multiple comparison test.

## Discussion

As a consequence of the COVID-19 pandemic, there has been a significant increase in research on coronavirus discovery, reservoir characterisation and spillover as well as the development of broad acting antivirals and therapeutics^21–24^. However, most studies have focused on betacoronaviruses, whereas the zoonotic and pandemic potential of alphacoronaviruses has remained relatively uncharacterised. Here we employed a new approach aimed at analysing and capitalising on viral heterogeneity at the genus wide level to gain a broad understanding of receptor usage and host tropism, since entry is the first key step during any viral spillover event. Using receptor libraries that included various mammalian species, we were able to classify the APN and ACE2 using viruses as generalists or specialists, based on their host range tropism. Broadly speaking, viruses were considered generalists if they could use the APN or ACE2 from various animal species to enter cells, whereas specialists were restricted to only a small number of receptors. Furthermore, we identified sub-classifications to these assignments (see Fig. 2): e.g. hCoV/NL63 was considered a ‘broad generalist’ since it could use the ACE2 receptors from various animals from different taxonomic orders to enter cells, whereas btCoV/AT1A-F41 was classified as an ‘order generalist’, since it could widely interact with the APN from various species, but only from a single order, in this case *Chiroptera*. With respect to specialists, hCoV/229E and btCoV/WA1087 were categorised as ‘multi-order specialists’, since they could use only a limited number of APNs from different animal orders to enter cells. Finally, the canine coronaviruses selected by our algorithm showed the most restricted host range, with these ‘true specialists’ only able to use the APN from canid species.

While our results established different classes of tropism amongst alphaCoVs using APN and ACE2, the broader trend was that use of these two receptors is poorly conserved across the genus. This was especially true for the 27 S proteins whose ORFs were sequenced following sampling of bats. Only 2/25 functionally pseudotyped bat virus S proteins used a recognised alphaCoV receptor. None of the true specialists characterised in our screens utilised receptors from only a single species (Fig. 2b), e.g. CCoVs A76 and SD-F3 could use APNs from other canids for entry (Fig. 6a). While we cannot formally discount the hypothesis that at least some bat alphaCoVs are hyper-specialised to only one cognate bat APN or ACE2, which was not included in our libraries, our results strongly suggest that one or more currently unknown receptors are involved in the life cycle of alphaCoVs.

Nevertheless, for those alphaCoVs that used a recognised receptor we were able to use structural data to demonstrate that the diversity of these phenotypic traits (generalism vs. specialiam) can be linked to individual amino acid residues (genotypes) within the S RBD-receptor interface. In fact, our structure-guided mutagenesis of hCoV/229E S showed that a single residue substitution, K316S, could substantially extend its host range to include several new bat species. Given the putative bat origin for 229E viruses, hypotheses which are based on the identification of 229E-related viruses in *Hipposideros* bats in Ghana^15^, the characterisation of this pivotal residue in host-range tropism provides insight into possible mechanisms of adaptation during spillover. This is especially true given the variability of this position in related sequences (Fig. 4b); all 229E-related S deposited in public databases possess a residue with a short side chain (S/V/T) at the respective 316 position, in contrast to K in human 229E. Another example of a single amino acid at the RBD-receptor interface being pivotal in determining host-range is ACE2 position 354, with glycine at this position being critical for NL63 entry (Fig. 3c). Despite its broad host range, no reports are available concerning the identification of NL63 in other mammals, and the evolutionary route that led to human adaptation is poorly understood. This might be due to absence of pathogenicity or clearance by host restriction factors, which would limit NL63 propagation and replication in hosts other than humans^25^. Our results show that phenotypic traits like receptor usage can vary significantly between closely related species. This variation can generally be explained by differences in protein-protein interactions at the RBD-receptor interface. This highlights the importance of detailed structural studies and mutagenesis to inform accurate prediction of the host range of emerging viruses.

The action of type II transmembrane serine proteases adds another dimension to the complexity of alphaCoV receptor tropism. The presence of catalytically-active human TMPRSS2 or HAT significantly expands the number of APN species that support btCoV/AT1A-F41 entry, from three to ten (Fig. 5a). This expansion, which to our knowledge is the first reported evidence for protease-dependent increases in host range, suggests that future investigations of viral receptor usage should consider the proteolytic status of the attachment and/or fusion proteins. We hypothesise that priming of S, which releases the fusion peptide, could favour rapid fusion upon binding to receptors that possess lower affinity, or that priming might affect the subcellular location where fusion takes place (e.g. plasma membrane versus endosome). Similarly, our comparisons of canine coronavirus host range provide significant insight into how receptor usage can vary within viral species. As discussed, the S proteins initially selected by our unbiased algorithmic method (CCoV/A76 and CCoV/SD-F3) behaved as true specialists in our receptor screening. However, the S proteins from CCoV associated with spillover into humans, had broad generalist phenotypes^26^ (Fig. 6a). Of note, these generalist CCoV isolates could not use human APN, which suggests entry into human cells may be conferred by another receptor. Indeed, previous studies have shown entry in cells that do not express APN, hinting at the role of other cellular constituents, such as lectins or sialic acids, in entry^26^. Our data also demonstrated that the generalist CCoV could tolerate swaps to its domains, whereas the RBD of the specialist CCoV seemed to be more functionally constrained compared with other domains in S. For example, CCoV/SD-F3 S was not functional when its loop 1 was replaced. These results suggests that attachment proteins could also be studied for their tolerance for recombination and intragenic stability, events which could facilitate the evolution of alphaCoV S towards generalism and an increased risk of spillover.

Evolution-driven variation of receptors across the animal kingdom, together with the overall affinity of the attachment protein-receptor virus-host interaction, likely plays a key role in determining the host-range of viruses. For example, NL63, like many ACE2 using sarbecoviruses, was found to be a broad generalist, while all APN-using alphaCoVs had more restricted tropisms. Nevertheless, we found that all the APN- or ACE2-using alphaCoV S included in our study could use the receptor of at least one peridomestic animal. The most interesting example is domestic cat APN: it supports entry for 229E-related viruses with great efficiency and is the only receptor not affected by protease co-expression. Flow cytometry analysis confirms that the broad usage of cat APN is not a consequence of higher transfection efficiency or APN expression in the transfected cells used for pseudotype infection assays (SFig. 4). This suggests that the molecular basis of the interaction between alphaCoV RBDs and cat APN differs from interactions with other APNs. In the future, it would be interesting to further investigate the mechanisms underlying this species-specific permissiveness, especially since domestic cats are known to hunt bats and could potentially act as an important intermediate reservoir for spillover^27^. We are currently examining other feline APN sequences to understand if this is phenotypically restricted to only domesticated cat species. Another important example of the divergent tropism of bat CoVs is btCoV/WA1087 which could efficiently use caprid (sheep and goat) APNs for entry. This is again relevant from the perspective of spillover preparedness and understanding the origin of zoonotic viruses. Further studies could be designed to assess if the virus could infect and replicate in these animals, especially considering the significant overlap in the host-range of Western Australian microbats, where btCoV/WA1087 was isolated, and small ruminant agriculture^28^. Additionally, a better understanding of this virus’s biology should be acquired regarding potential recombinant with closely related porcine coronaviruses, especially in the context of small farms where different animal species are kept together. In fact, btCoV/WA1087 as a pedacovirus, contains recombination hotspots in the genome^29^ and includes economically relevant porcine viruses PEDV and PRCV. Lastly, although little is known about the canine coronaviruses that have crossed the species barrier and potentially infected humans^30^, our data suggest that an intermediate host cannot be excluded in the process of transmission to humans.

The ‘holy grail’ of pandemic preparedness is to be able to predict the zoonotic potential of viruses from their genome sequence alone. Various approaches are being leveraged to do this, one such being genotypic-phenotypic linkage of individual traits and the application of machine learning algorithms to predict zoonotic potential^31^. Our hybrid approach merges the advantage of unbiased selection of informative sequences from data-rich resources, such as Genbank, with experimental data and high-throughput screenings. Our data highlights that phenotypic traits such as host-range can vary greatly, even within single species. Predictions of spillover potential, especially when using *in silico* approaches, must reflect this in their design in order to maximise their power. However, it may be that the ‘inherent unpredictability’ of receptor usage as a trait will improve predictive analysis when combined with other more conserved aspects of the viral life cycle, e.g. innate immune modulation, replication etc. To move forward with computational predictions and more generally to improve our understanding of how viral entry determines spillover potential, additional experimental data, such as targeted or deep mutational scanning mutagenesis, should be obtained. Perhaps more importantly, it is worth stressing that negative results are as informative as positive, in this case highlighting that our knowledge of alphaCoV receptor(s) usage is relatively limited.

## Material and methods

### Construction of gene libraries

A schematic of the downstream analysis pipeline used for retrieving alphacoronavirus data and generating the final spike protein library used in this study (n=40) is provided in Figure 1a. All publicly available alphacoronavirus genome sequences were retrieved from the Virus Pathogen Database and Analysis Resource (ViPR) platform hosted by the Bioinformatics Resource Center (BRC) at the National Institute of Allergy and Infectious Diseases (NIAID)^32^. As of May 2021, the full database consisted of 19,082 alphacoronavirus genomes from which we extracted spike protein coding sequences, to a final database of 2,714 sequences. We constructed the spike proteins alignment using MAFFT 7.526^33, 34^ by integrating structural alignments of homologous spike protein structures queried from the UniProt Reference Clusters (UniRef)^35^. We then run maximum-likelihood phylogenetic reconstruction in IQTREE 2.3.4^36^ with 1000 ultrafast bootstrap replicates (UFBoot) and 1000 SH-like approximate likelihood ratio tests (aLRTs)^37^. Pairwise distances of branch lengths were computed between pair of tips using the *ape* package^38^ in R 4.4.1, which informed the unbiasing selection of 40 spike protein coding sequences by applying a greedy algorithm. Although heuristic, this selection approach ensures that an optimal subset of tips (i.e. evolutionary units) is returned under the assumption of maximising both minimum phylogenetic distance and phylogenetic diversity, as theoretically discussed by Bordewich et al.^39^. Genbank accession numbers of the S sequences used for the study are provided in Table S4.

### Plasmids used for pseudotyping

Selected alphaCoV S sequences were ordered from Biobasic (Canada) as codon-optimized synthetic genes, subcloned in pcDNA3.1 with an HA tag in the C-terminal cytoplasmic tail. For the APN library, genes were ordered from GenScript (UK) and subcloned with a N-terminal V5 tag in pCAGGS vector. For the ACE2 library and human DPP4 (Genbank: NP_001926), the ectodomains of the ORFs were subcloned with a N-terminal HA tag in pDisplay. Untagged human TMPRSS2 (Genbank: NP_001128571.1) and its mutated form S441A, along with human HAT, were cloned in pcDNA3.1. Genbank accession numbers of the sequences used for the study are provided in Table S2 and S3 for ACE2 and APN libraries, respectively.

### Cells

Human renal HEK293T were cultured in Dulbecco’s Modified Eagle Medium (DMEM) supplemented with 10% Fetal Bovine Serum (FBS), penicillin-streptomycin (PenStrep) and sodium pyruvate. All reagents for cell culture were purchased from Gibco.

### Pseudotype virus production

To pseudotype alphaCoV S, plasmids encoding their ORF were transfected in confluent HEK293T cells seeded in a 6 well plate using polyethylenimine (PEI, 5 µg/mL). HIV-1-based lentiviral vectors coding for viral structural proteins were also included in the transfection mix. The following day, media was replaced, and pseudoviruses in the supernatant harvested on 48 and 72 h post-transfection and pooled together. The last day, supernatant was spun down 10 min at 4000 rpm, to remove cellular debris. Finally, pseudoviruses were aliquoted and stored at -80°C, until further use. To verify S incorporation, pseudoparticles were purified by ultracentrifugation at 23,000 rpm, 4°C for 2 h using a 20% sucrose gradient. Supernatants were discarded and pellet resuspended in PBS. Concentrated pseudoparticles were lysed by boiling with Laemmli (Biorad) and S expression analysed by sodium dodecyl sulfate polyacrylamide gel electrophoresis (SDS-PAGE). Separated proteins were transferred on a 0.45 µm nitrocellulose membrane (Cytiva), blocked in PBS supplemented with 0.05% TWEEN-20 and 5% (w/v) unskimmed milk powder, and incubated with mouse monoclonal anti-HA (clone 6E2, Cell Signalling Technology) and anti-p24 (clone 5, Abcam) overnight at 4°C. The following day, goat anti-mouse DyLight 680 (Invitrogen) was used to probe primary antibodies, and signals detected with an Odyssey DLx imaging system (LI-COR Biosciences).

### Pseudovirus entry assay

For receptor usage screening, HEK293T were transfected with plasmids coding for the receptors of interested. The following day, media was removed and replaced with fresh DMEM supplemented with 10% FBS, to have a final concentration of 2×10^4^ cells/mL. 100 µL of cell preparation were aliquot in each well of a 96 well plate, and treated with 100 µL of pseudovirus preparation, diluted 1:1 with fresh media. Two days later, supernatant was removed and cells were treated with Bright-Glo (Promega), diluted 1:1 with PBS. Luciferase signals were acquired using a Glomax Discover luminometer (Promega).

To assess cell line permissivity to pseudovirus infection, confluent human cell lines were treated with undiluted pseudoparticle supernatant in a 96 well plate. Two days later, plates were spun down, media replaced with 50 µL of Bright-Glo and luciferase signals acquired on the luminometer.

### Flow cytometry for expression of the receptor libraries

Plasmids were transfected in HEK293T using Trans-IT X2 (Mirus Bio). The following day, cells were resuspended in PBS, fixed using 2% paraformaldehyde for 20 min at 4°C and permeabilized using PBS supplemented with 0.5% Triton X100 for 5 min at 4°C. After washing, cells were incubated for 1 h with an anti-tag antibody conjugated with PE and APC for APN and ACE2 libraries, respectively. Samples were analysed using a MACSQuant Analyzer 10 cytometer (Miltenyi Biotec), and data analysis performed using FlowJo (BD Biosciences).

### Site-directed mutagenesis

Substitutions of S amino acids were introduced using Quikchange Lightning Site-Directed Mutagenesis kit (Agilent) following manufacturer’s instructions. Primers were designed using Agilent online tool (https://www.agilent.com/store/primerDesignProgram.jsp). To swap proteins’ domains between different CCoV strain, their RBDs or RBD loops were amplified by PCR and inserted in the plasmid using HiFi Assembly (NEB).

### Computational analysis

Figures of structural models were rendered using PyMOL (Schrödinger). Sequence conservation was obtained as pdb models on deposited structures using ESPript3.0^40^ by supplying the alignments of APN or RBD sequences. GraphPad Prism 9 was used to obtain all the other figures, and to perform statistical analysis.

## Acknowledgements

We would like to thank Thomas Peacock, Nazia Thakur and Joe Newman at The Pirbright Institute for provision of the TMPRSS2 plasmis, other protease constructs, and the ACE2 expression library. In addition, we would like to thank all coronavirus research teams that deposited sequence information to Genbank that were selected by our greedy-algorithm based approach (accession numbers available in Table S1).

## References

1. Ellwanger, J.H. & Chies, J.A.B. Zoonotic spillover: Understanding basic aspects for better prevention. Genet Mol Biol 44, e20200355 (2021).

2. Tan, C.C.S., van Dorp, L. & Balloux, F. The evolutionary drivers and correlates of viral host jumps. Nat Ecol Evol 8, 960–971 (2024).

3. Iwata-Yoshikawa, N. et al. TMPRSS2 Contributes to Virus Spread and Immunopathology in the Airways of Murine Models after Coronavirus Infection. J Virol 93 (2019).

4. Yeager, C.L. et al. Human aminopeptidase N is a receptor for human coronavirus 229E. Nature 357, 420–422 (1992).

5k5 . Li, Z. et al. The human coronavirus HCoV-229E S-protein structure and receptor binding. Elife 8 (2019).

6. Reguera, J. et al. Structural bases of coronavirus attachment to host aminopeptidase N and its inhibition by neutralizing antibodies. PLoS Pathog 8, e1002859 (2012).

7. Tusell, S.M., Schittone, S.A. & Holmes, K.V. Mutational analysis of aminopeptidase N, a receptor for several group 1 coronaviruses, identifies key determinants of viral host range. J Virol 81, 1261–1273 (2007).

8. Hofmann, H. et al. Human coronavirus NL63 employs the severe acute respiratory syndrome coronavirus receptor for cellular entry. Proc Natl Acad Sci U S A 102, 7988–7993 (2005).

9. Cui, X. et al. Virus diversity, wildlife-domestic animal circulation and potential zoonotic viruses of small mammals, pangolins and zoo animals.

10. Wang, D. et al. Substantial viral diversity in bats and rodents from East Africa: insights into evolution, recombination, and cocirculation.

11. Tao, Y. et al. Surveillance of Bat Coronaviruses in Kenya Identifies Relatives of Human Coronaviruses NL63 and 229E and Their Recombination History. J Virol 91 (2017).

12. Frank, H.K., Enard, D. & Boyd, S.D. Exceptional diversity and selection pressure on coronavirus host receptors in bats compared to other mammals. Proc Biol Sci 289, 20220193 (2022).

13. Liu, C. et al. Receptor usage and cell entry of porcine epidemic diarrhea coronavirus. J Virol 89, 6121–6125 (2015).

14. Ntumvi, N.F. et al. Wildlife in Cameroon harbor diverse coronaviruses, including many closely related to human coronavirus 229E. Virus Evol 8, veab110 (2022).

15. Corman, V.M. et al. Evidence for an Ancestral Association of Human Coronavirus 229E with Bats. J Virol 89, 11858–11870 (2015).

16. Hulswit, R.J., de Haan, C.A. & Bosch, B.J. Coronavirus Spike Protein and Tropism Changes. Adv Virus Res 96, 29–57 (2016).

17. Hoffmann, M., Hofmann-Winkler, H. & Pöhlmann, S. Priming Time: How Cellular Proteases Arm Coronavirus Spike Proteins.

18. Escalante, D.E. & Ferguson, D.M. Structural modeling and analysis of the SARS-CoV-2 cell entry inhibitor camostat bound to the trypsin-like protease TMPRSS2. Med Chem Res 30, 399–409 (2021).

19. Vlasova, A.N. et al. Novel Canine Coronavirus Isolated from a Hospitalized Patient With Pneumonia in East Malaysia. Clin Infect Dis 74, 446–454 (2022).

20. Lednicky, J.A. et al. Isolation of a Novel Recombinant Canine Coronavirus From a Visitor to Haiti: Further Evidence of Transmission of Coronaviruses of Zoonotic Origin to Humans. Clin Infect Dis 75, e1184–e1187 (2022).

21. Kuchinski, K.S. et al. Targeted genomic sequencing with probe capture for discovery and surveillance of coronaviruses in bats. Elife 11 (2022).

22. Ghai, R.R. et al. Animal Reservoirs and Hosts for Emerging Alphacoronaviruses and Betacoronaviruses. Emerg Infect Dis 27, 1015–1022 (2021).

23. Karim, M., Lo, C.W. & Einav, S. Preparing for the next viral threat with broad-spectrum antivirals. J Clin Invest 133 (2023).

24. Monrad, J.T., Sandbrink, J.B. & Cherian, N.G. Promoting versatile vaccine development for emerging pandemics. NPJ Vaccines 6, 26 (2021).

25. Li, Q. et al. Cross-species transmission, evolution and zoonotic potential of coronaviruses. Front Cell Infect Microbiol 12, 1081370 (2022).

26. Liu, Y. et al. Characterization of CCoV-HuPn-2018 spike protein-mediated viral entry. J Virol 97, e0060123 (2023).

27. Szentivanyi, T., Oedin, M. & Rocha, R. Cat–wildlife interactions and zoonotic disease risk: a call for more and better community science data. Mammal Review 54, 93–104 (2024).

28. Prada, D., Boyd, V., Baker, M.L., O’Dea, M. & Jackson, B. Viral Diversity of Microbats within the South West Botanical Province of Western Australia. Viruses 11, 1157 (2019).

29. Han, Y. et al. Panoramic analysis of coronaviruses carried by representative bat species in Southern China to better understand the coronavirus sphere. Nat Commun 14, 5537 (2023).

30. Vlasova, A.N. et al. Animal alphacoronaviruses found in human patients with acute respiratory illness in different countries. Emerg Microbes Infect 11, 699–702 (2022).

31. Wardeh, M., Blagrove, M.S.C., Sharkey, K.J. & Baylis, M. Divide-and-conquer: machine-learning integrates mammalian and viral traits with network features to predict virus-mammal associations. Nat Commun 12, 3954 (2021).

32. Olson, R.D. et al. Introducing the Bacterial and Viral Bioinformatics Resource Center (BV-BRC): a resource combining PATRIC, IRD and ViPR. Nucleic Acids Res 51, D678–d689 (2023).

33. Katoh, K. & Standley, D.M. MAFFT multiple sequence alignment software version 7: improvements in performance and usability. Mol Biol Evol 30, 772–780 (2013).

34. Rozewicki, J., Li, S., Amada, K.M., Standley, D.M. & Katoh, K. MAFFT-DASH: integrated protein sequence and structural alignment. Nucleic Acids Res 47, W5–w10 (2019).

35. Suzek, B.E., Wang, Y., Huang, H., McGarvey, P.B. & Wu, C.H. UniRef clusters: a comprehensive and scalable alternative for improving sequence similarity searches. Bioinformatics 31, 926–932 (2015).

36. Minh, B.Q. et al. IQ-TREE 2: New Models and Efficient Methods for Phylogenetic Inference in the Genomic Era. Mol Biol Evol 37, 1530–1534 (2020).

37. Hoang, D.T., Chernomor, O., von Haeseler, A., Minh, B.Q. & Vinh, L.S. UFBoot2: Improving the Ultrafast Bootstrap Approximation. Mol Biol Evol 35, 518–522 (2018).

38. Paradis, E. & Schliep, K. ape 5.0: an environment for modern phylogenetics and evolutionary analyses in R. Bioinformatics 35, 526–528 (2019).

39. Bordewich, M., Rodrigo, A.G. & Semple, C. Selecting taxa to save or sequence: desirable criteria and a greedy solution. Syst Biol 57, 825–834 (2008).

40. Robert, X. & Gouet, P. Deciphering key features in protein structures with the new ENDscript server. Nucleic Acids Res 42, W320–324 (2014).

41. Li, F., Li, W., Farzan, M. & Harrison, S.C. Structure of SARS coronavirus spike receptor-binding domain complexed with receptor. Science 309, 1864–1868 (2005).

42. Lan, J. et al. Structure of the SARS-CoV-2 spike receptor-binding domain bound to the ACE2 receptor. Nature 581, 215–220 (2020).

43. Lee, J. et al. Broad receptor tropism and immunogenicity of a clade 3 sarbecovirus. Cell Host Microbe 31, 1961–1973.e1911 (2023).

44. Xiong, Q. et al. Close relatives of MERS-CoV in bats use ACE2 as their functional receptors. Nature 612, 748–757 (2022).

45. Wu, K., Li, W., Peng, G. & Li, F. Crystal structure of NL63 respiratory coronavirus receptor-binding domain complexed with its human receptor. Proc Natl Acad Sci U S A 106, 19970–19974 (2009).

46. Wong, A.H.M. et al. Receptor-binding loops in alphacoronavirus adaptation and evolution. Nat Commun 8, 1735 (2017).

47. Tortorici, M.A. et al. Structure, receptor recognition, and antigenicity of the human coronavirus CCoV-HuPn-2018 spike glycoprotein. Cell 185, 2279–2291.e2217 (2022).

48. Ji, W. et al. Structures of a deltacoronavirus spike protein bound to porcine and human receptors. Nat Commun 13, 1467 (2022).

