## Supplementary information for "Characterising alphacoronavirus phenotypic traits through diversity-driven selection of spike"

Fig 1 - Spikes and receptors' libraries information and RBD-receptors binding interfaces.

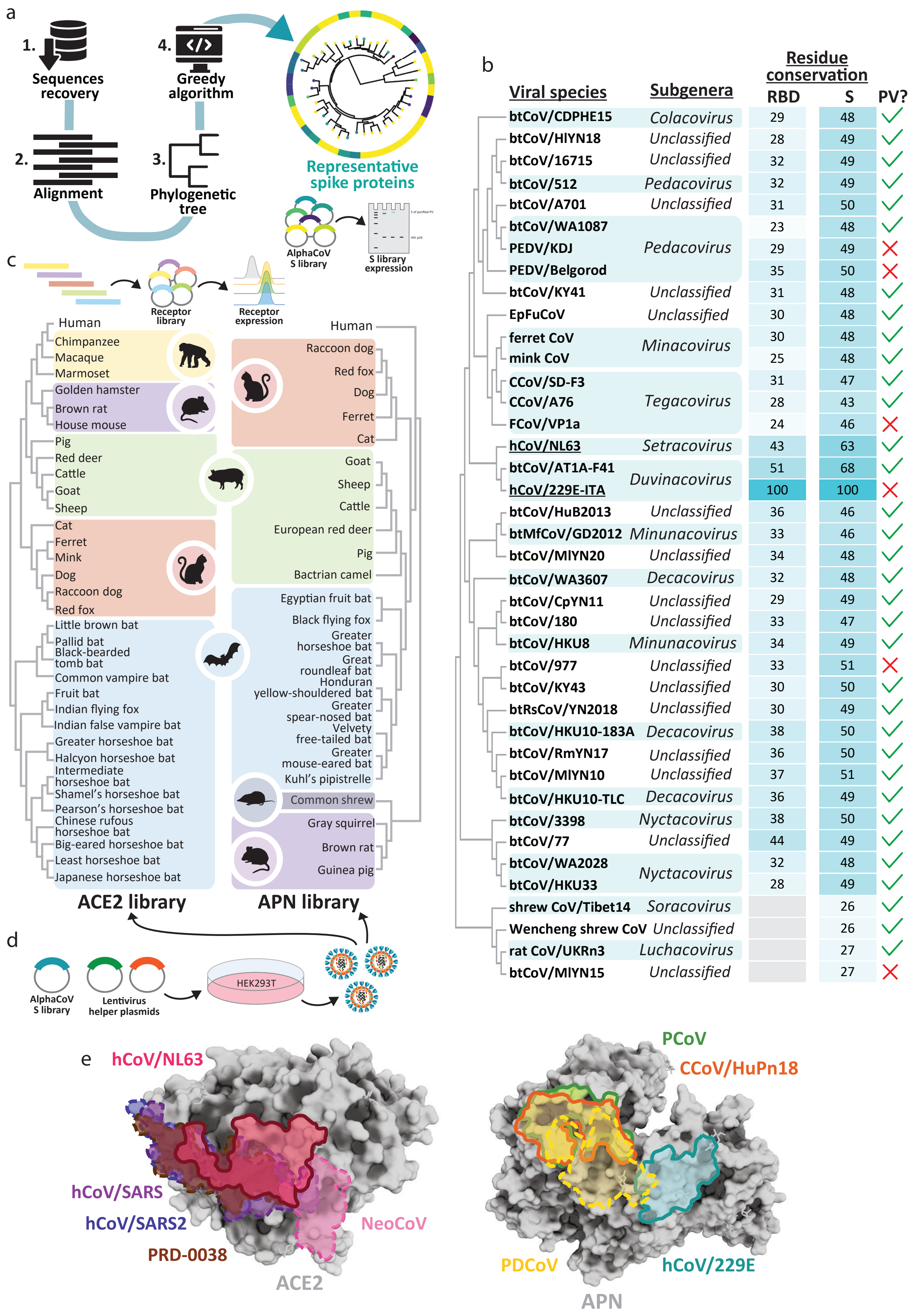

Fig 2 - AlphaCoV S usage of ACE2 and APN libraries, in absence or presence of cellular transmembrane protease TMPRSS2.

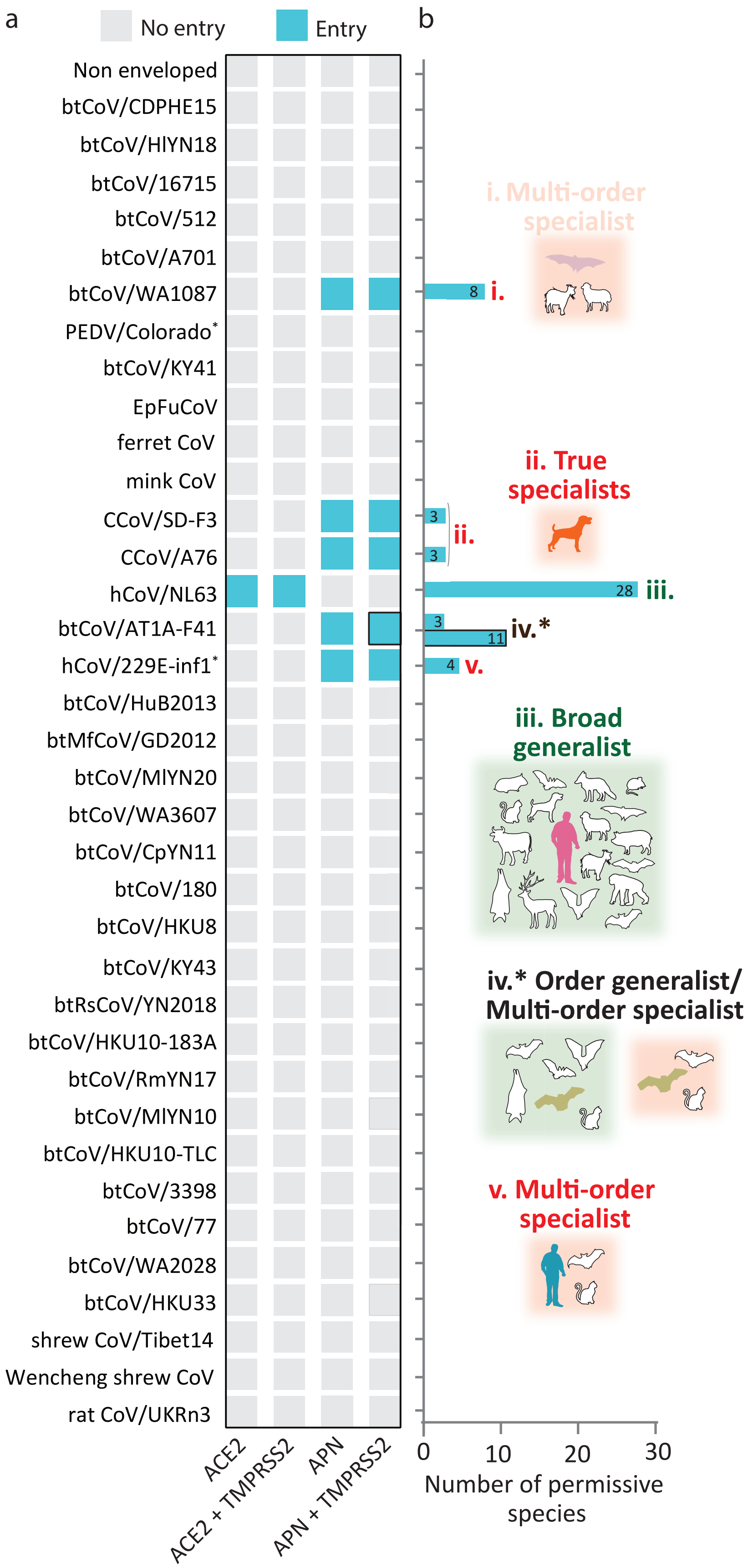

Fig 3 - Animal host range of human alphaCoVs NL63 and 229E S with their respective receptor, ACE2 and APN, and their molecular determinants of interactions

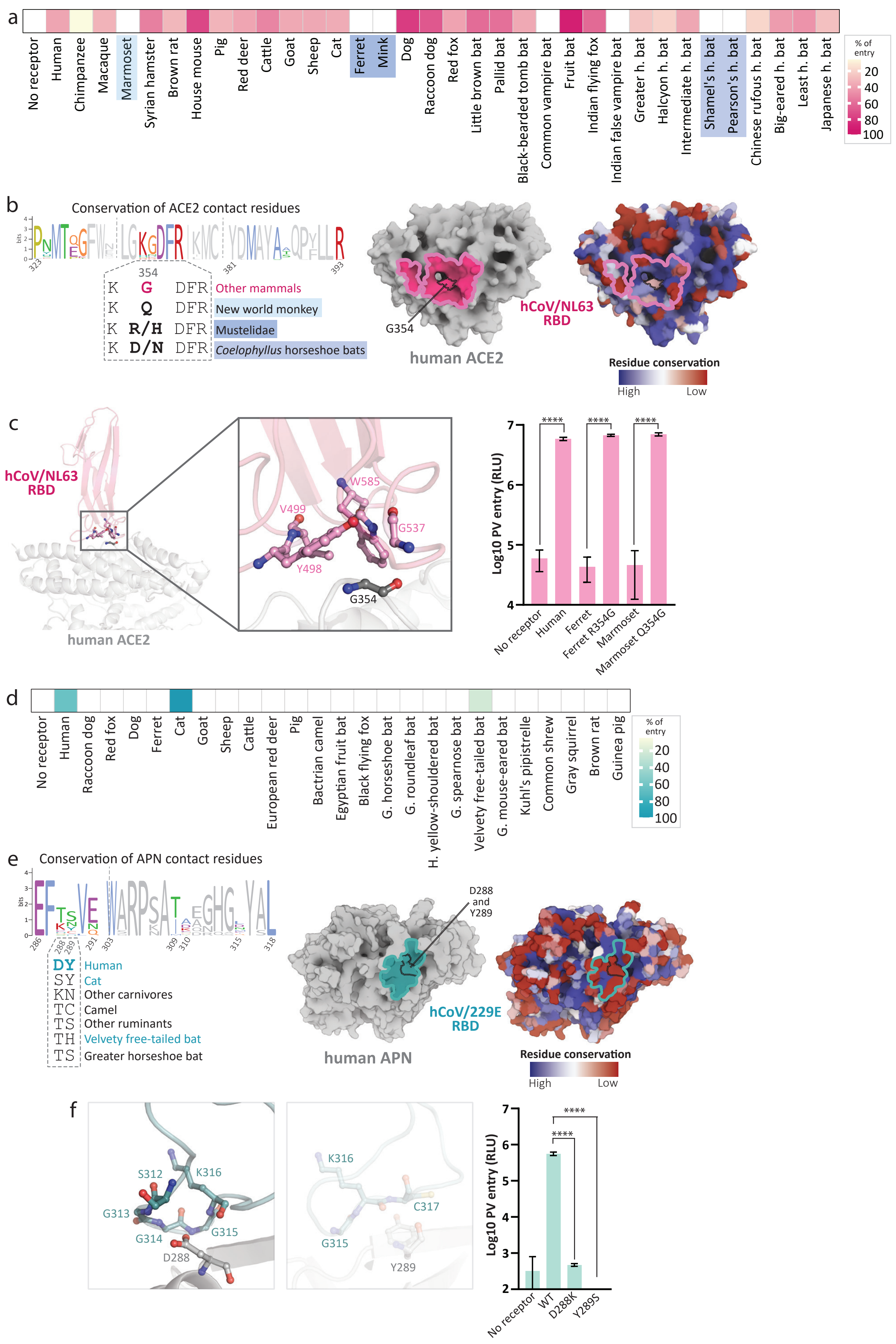

Fig 4 - Animal host range of 229E-like bat alphaCoV AT1A-F41, and the impact of substitions on 229E viruses on host range

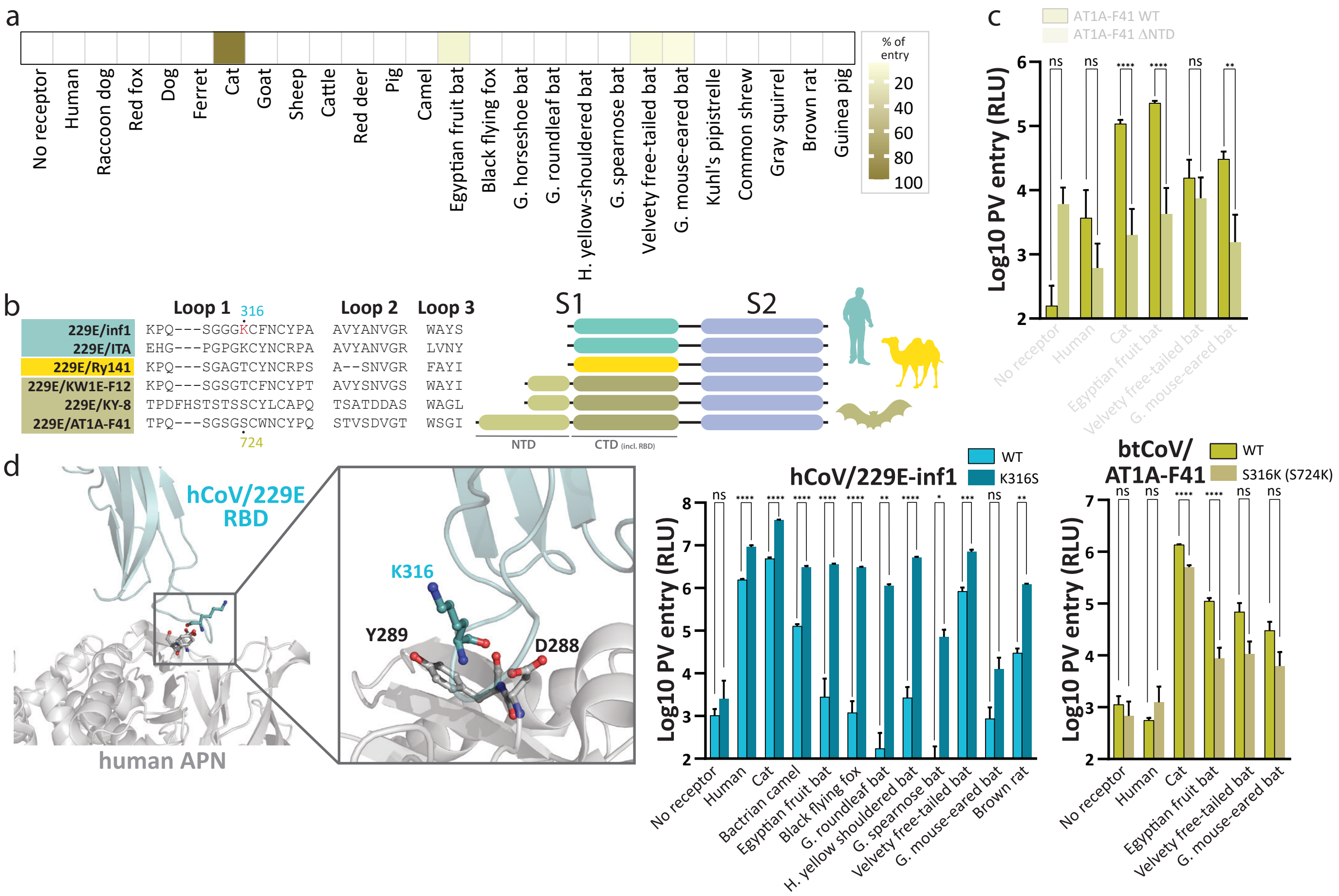

Fig 5 - Host range of WA1087, an alphaCoV using APN, and the impact of cellular proteases priming on bat-derived alphaCoV S using APN

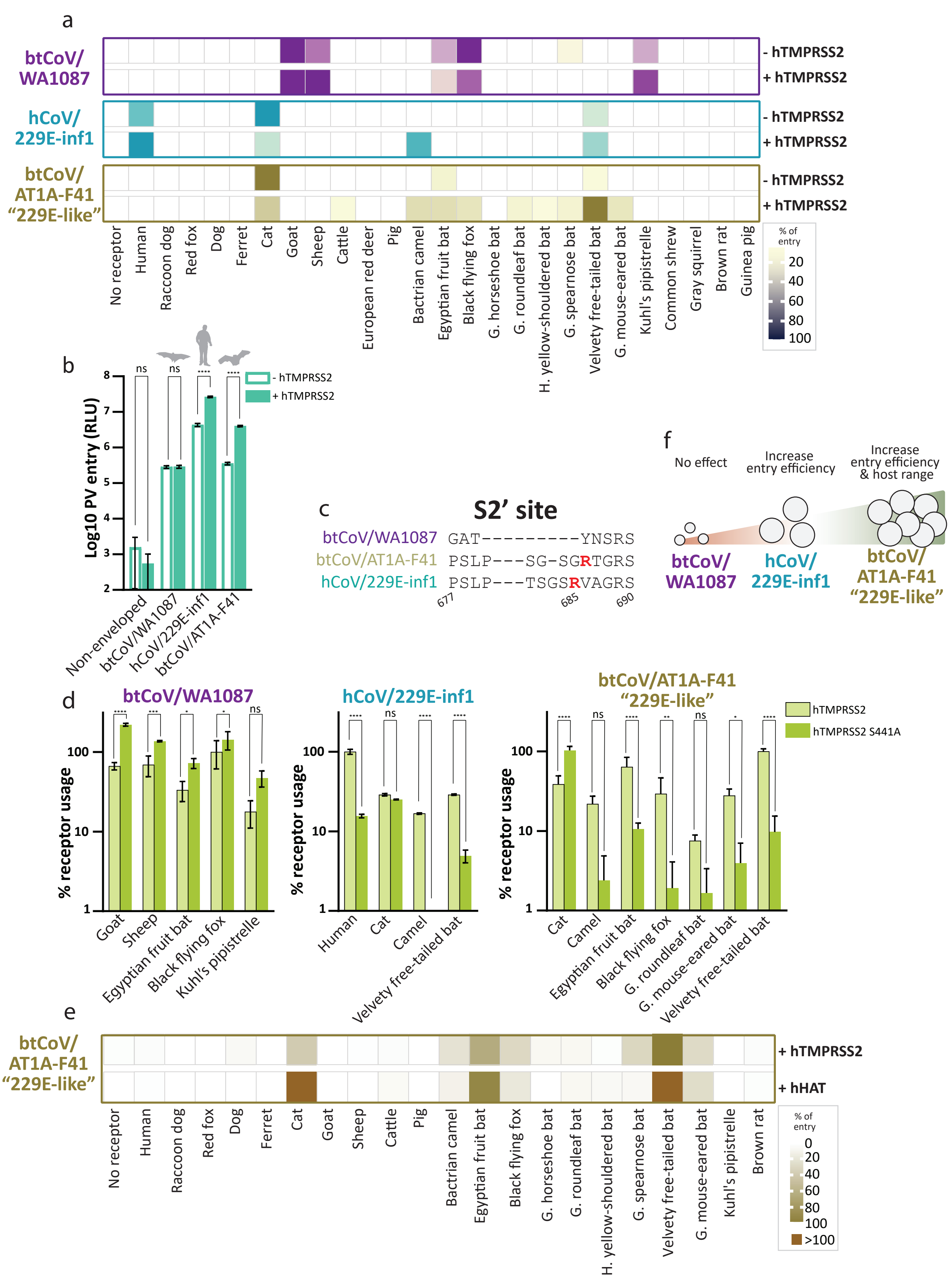

Fig 6 - Animal host range of canine alphaCoVs and the contribution of their RBD, or its subdomains, to APN usage

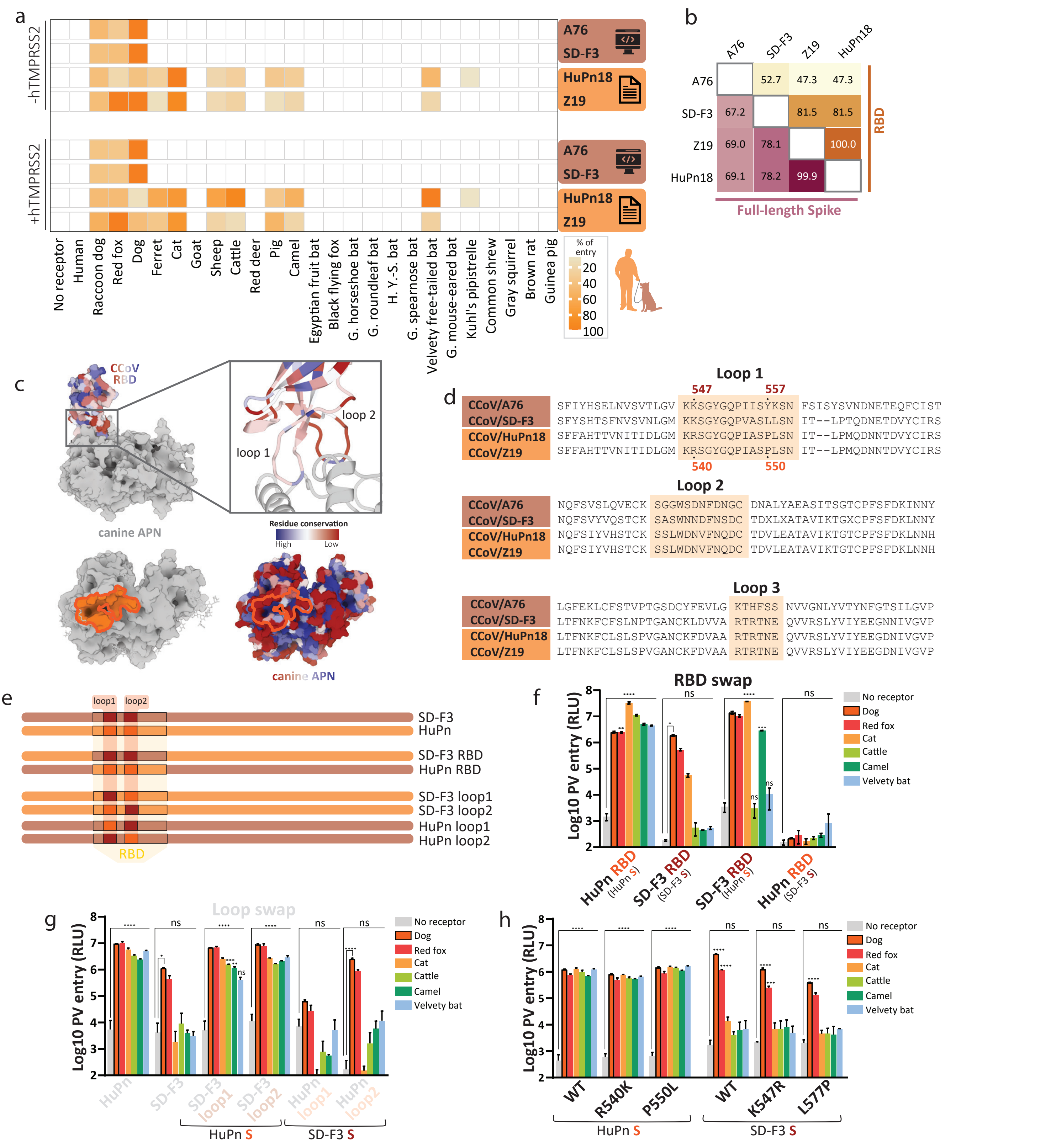

Fig S1 - Charateristics of alphaCoV Spikes included in the built library.

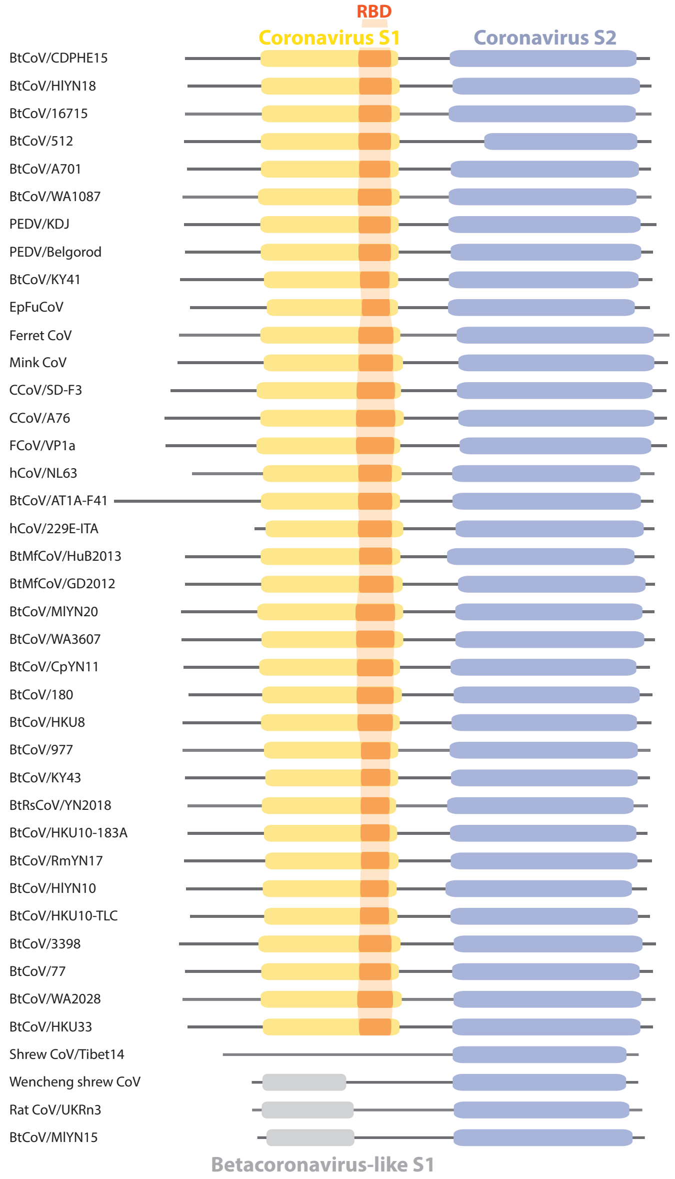


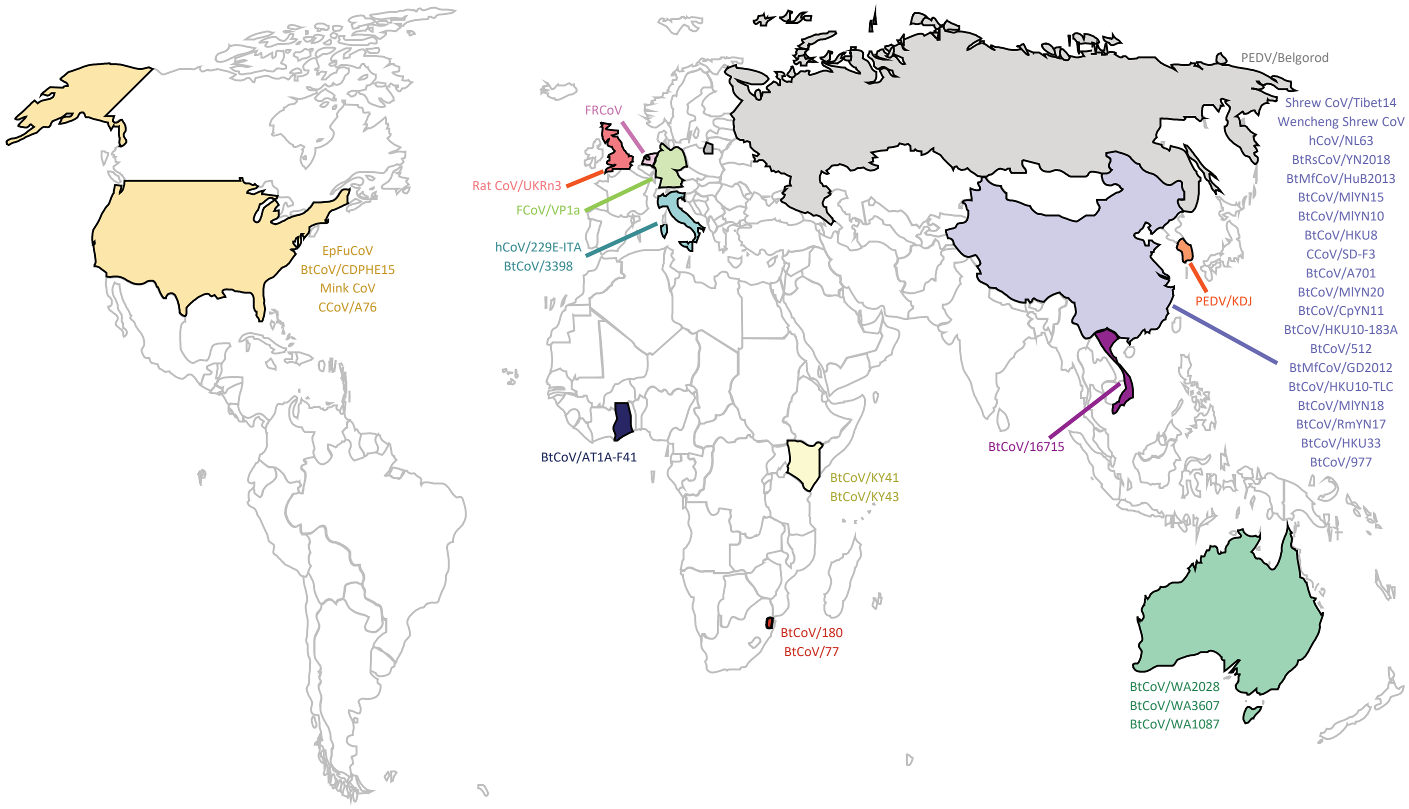

Fig S2 - Presentation of the alphaCoV S on the surface of pseudoparticles.

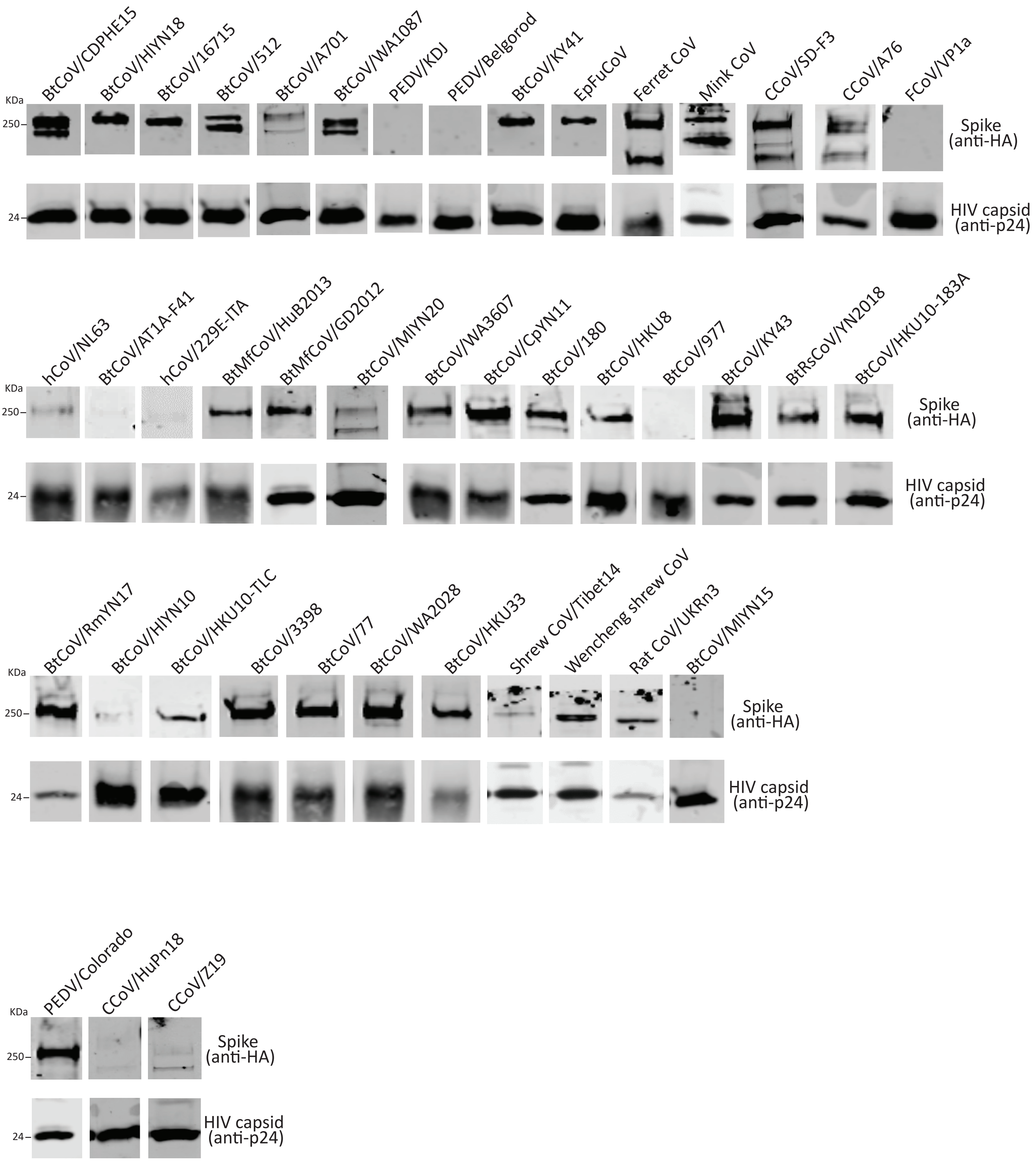

Fig S3 - Pipeline of S library study

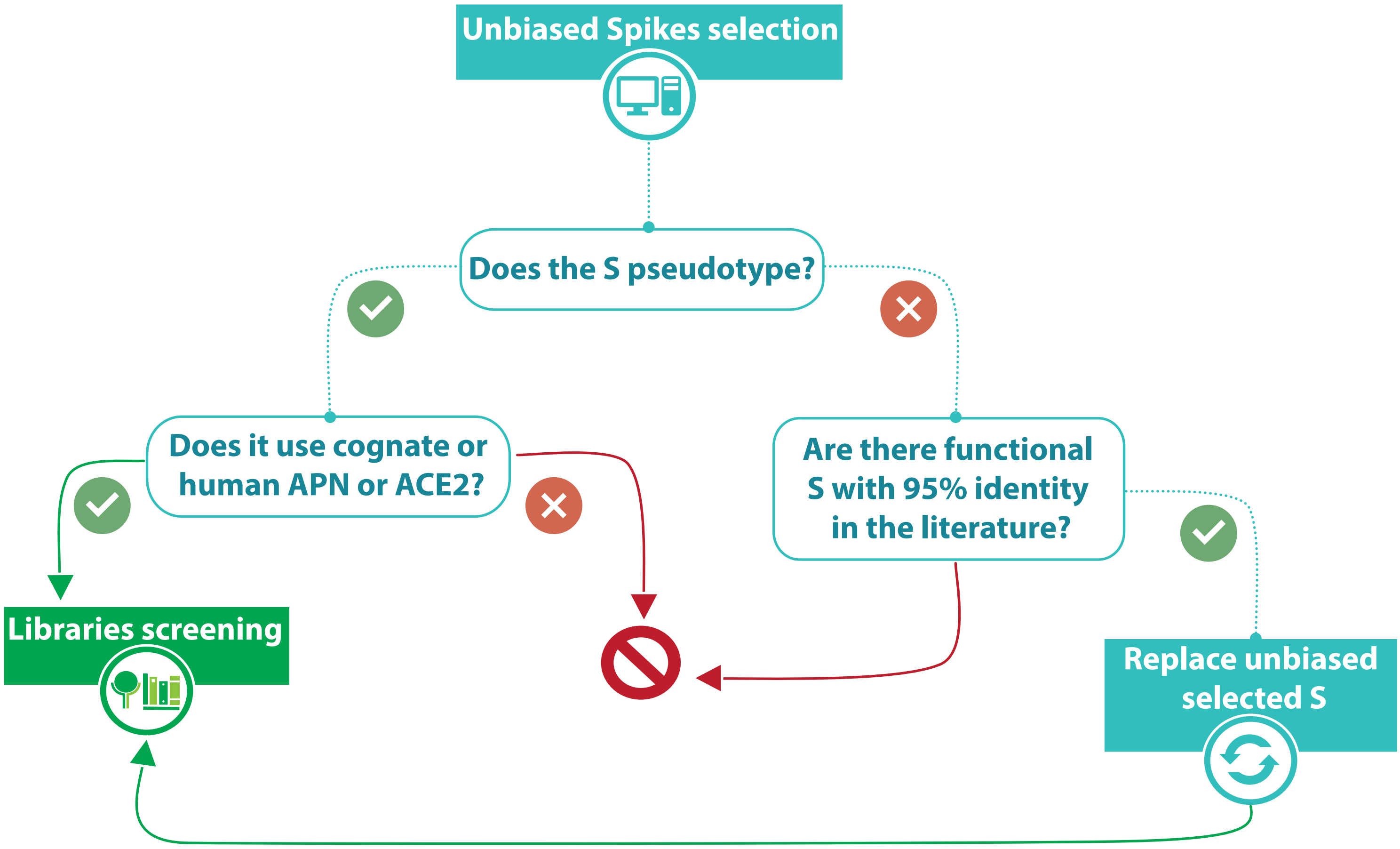

Fig S4 - Expression of receptors' libraries

a

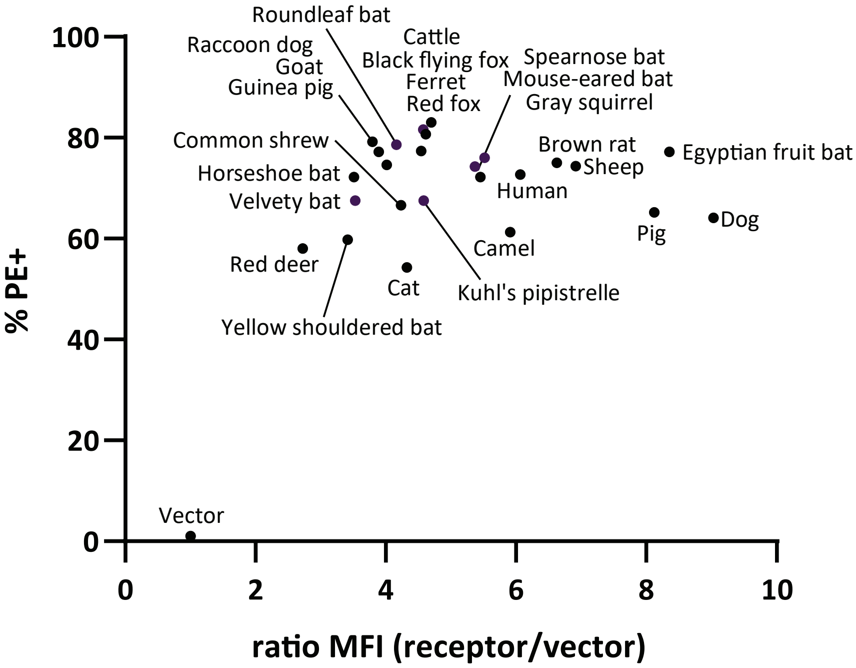


b

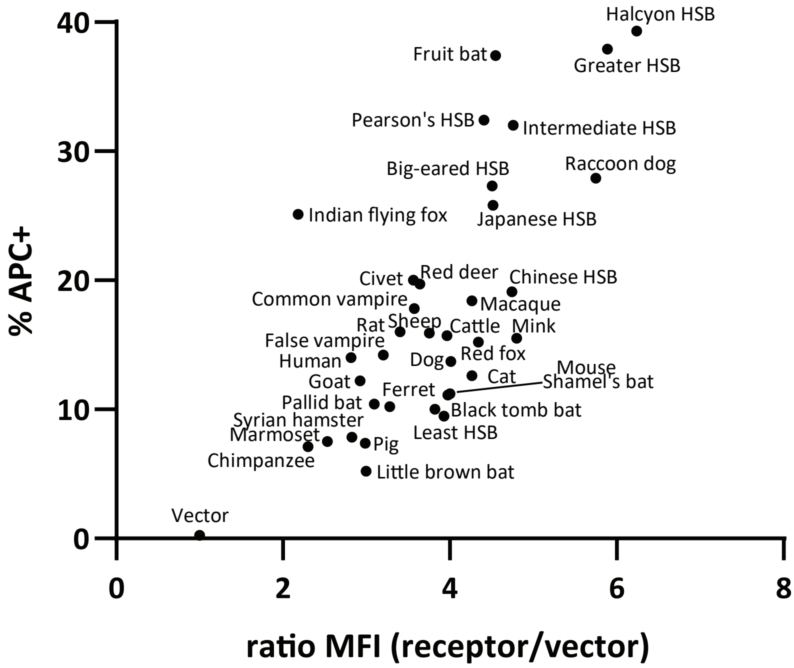

### Fig S5 - APN screening of alphaCoV library

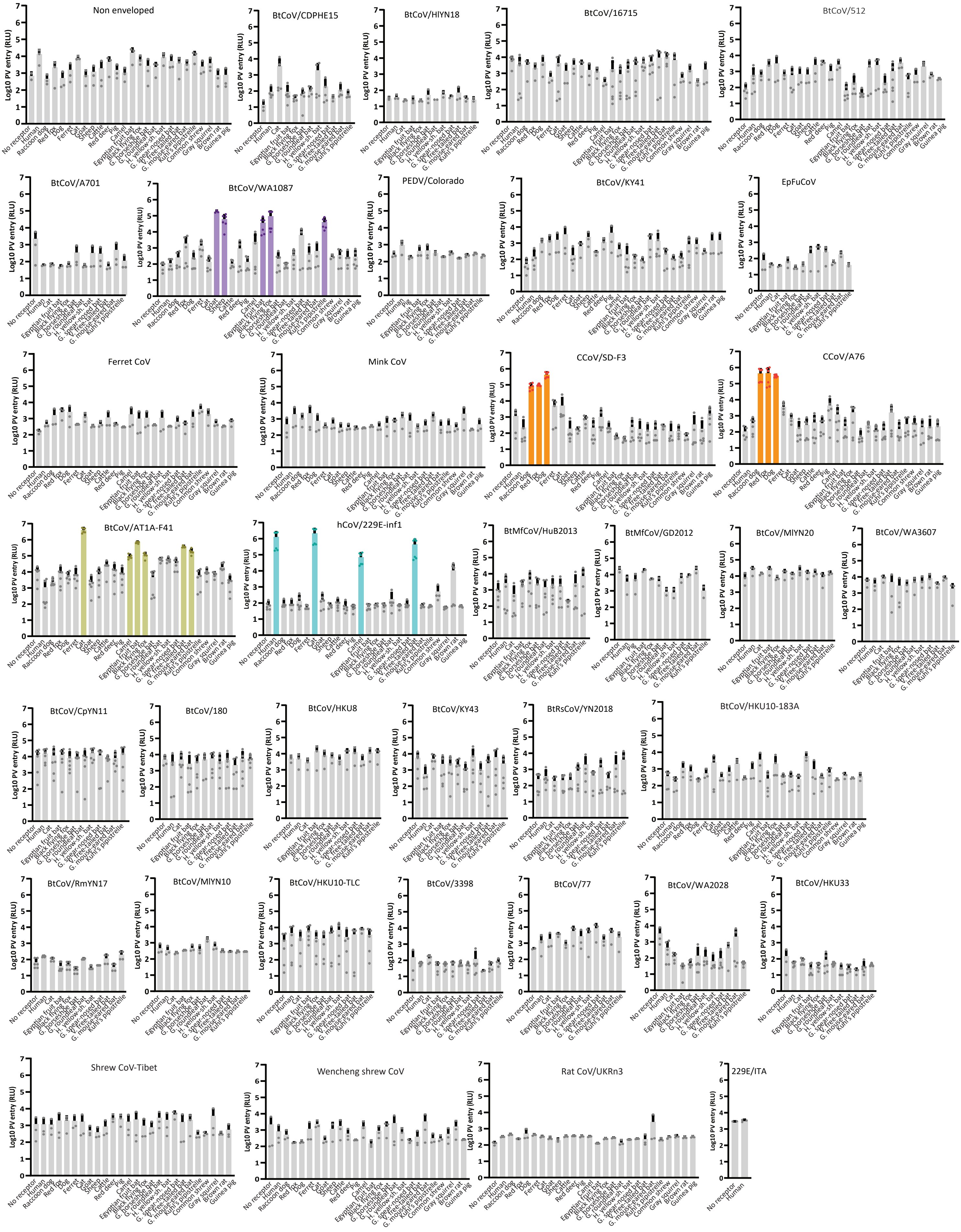

### Fig S6 - ACE2 screening of alphaCoV library

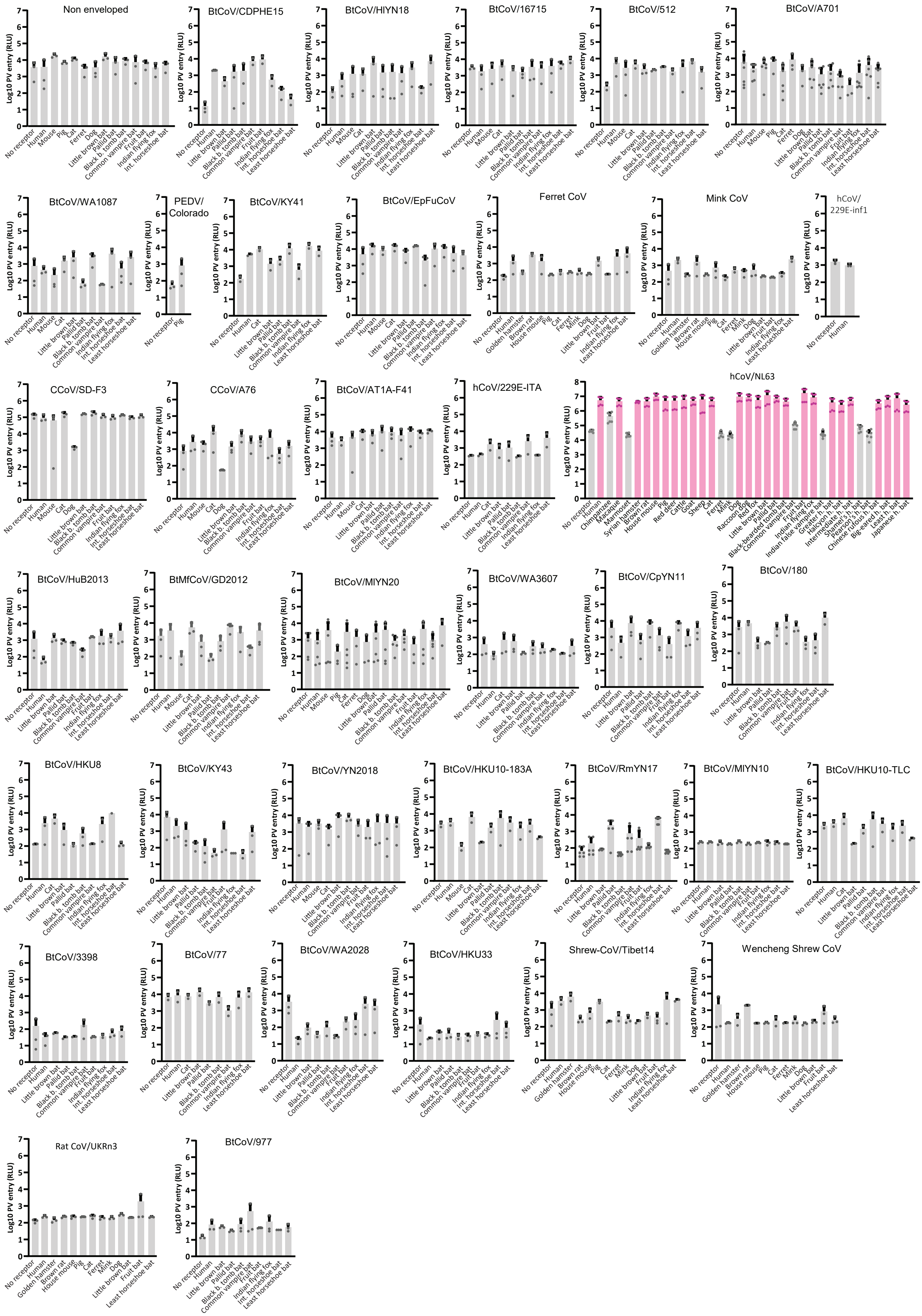

### Fig S7 - APN screening of alphaCoV library in presence of human TMPRSS2

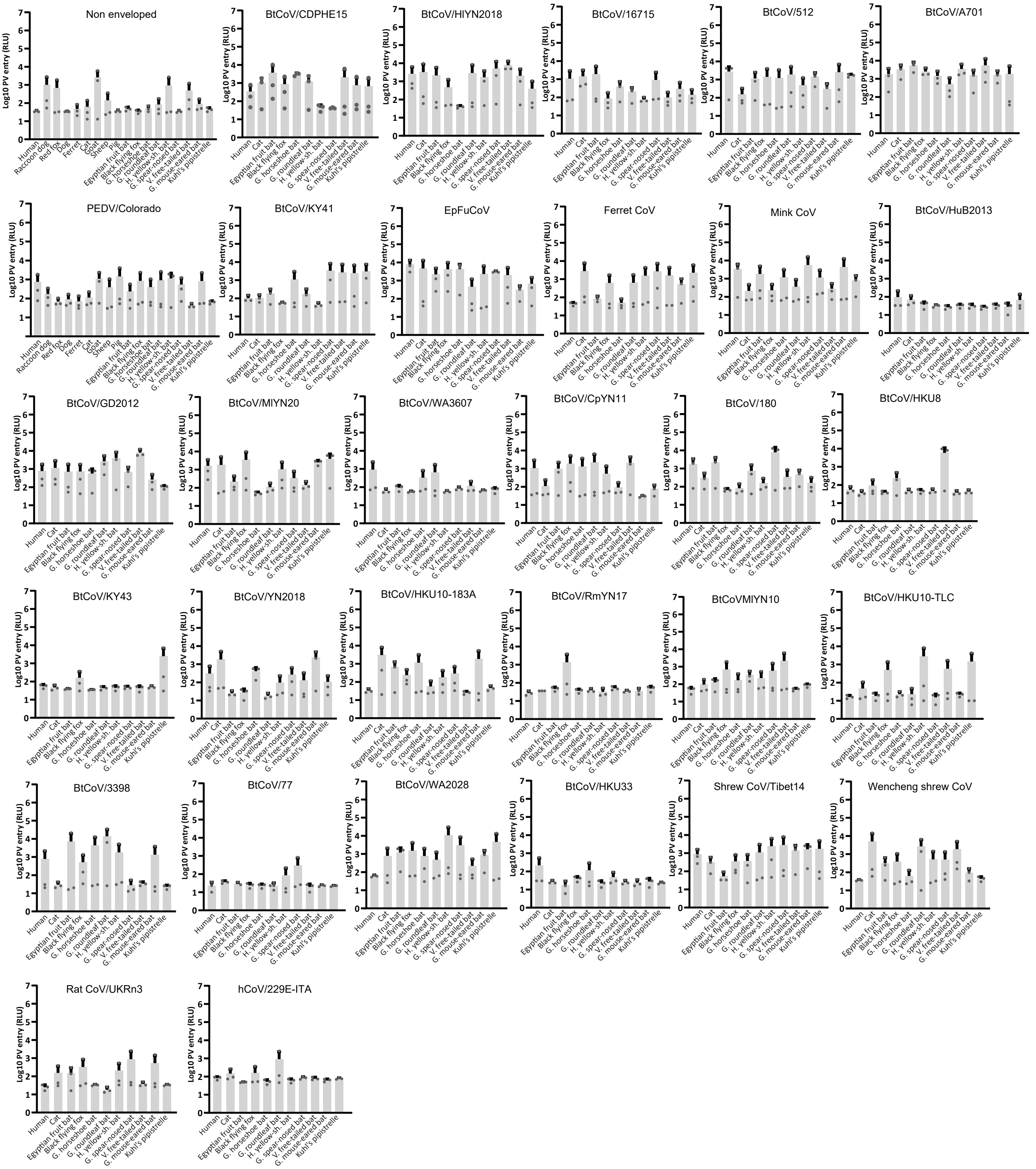

### Fig S8 - ACE2 screening of alphaCoV library in presence of human TMPRSS2

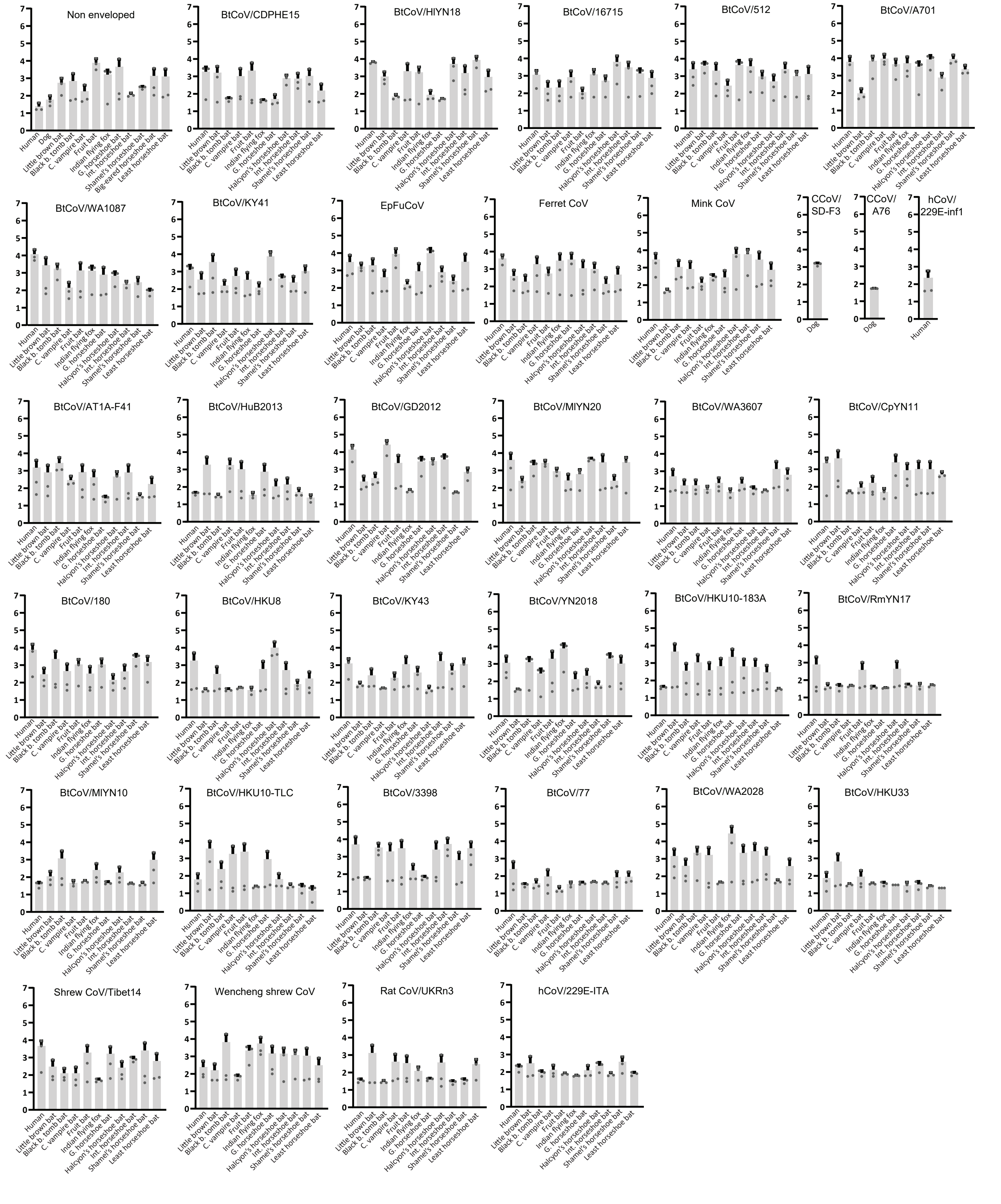

Fig S9 - Host range of APN-tropic alphaCoV included in the study

| | btCoV/ WA1087 | CCoV/ SD-F3 | CCoV/ A76 | btCoV/ AT1A-F41 | hCoV/ 229E |
|---|---|---|---|---|---|
| Human | - | - | - | - | |
| Raccoon dog | - | | | - | - |
| Red fox | - | | | - | - |
| Dog | - | | | - | - |
| Ferret | - | - | - | - | - |
| Cat | - | - | - | | |
| Goat | | - | - | - | - |
| Sheep | | - | - | - | - |
| Cattle | - | - | - | | - |
| European red deer | - | - | - | - | - |
| Pig | - | - | - | - | - |
| Bactrian camel | - | - | - | | |
| Egyptian fruit bat | | - | - | | - |
| Black flying fox | | - | - | | - |
| G. horseshoe bat | - | - | - | - | - |
| G. roundleaf bat | - | - | - | | - |
| H. yellow-shouldered bat | - | - | - | | - |
| G. spear-nose bat | | - | - | | - |
| Velvety free-tailed bat | - | - | - | | |
| G. mouse-eared bat | - | - | - | | - |
| Kuhl's pipistrelle | | - | - | - | - |
| Common shrew | - | - | - | - | - |
| Gray squirrel | - | - | - | - | - |
| Brown rat | - | - | - | - | - |
| Guinea pig | - | - | - | - | - |

Fig S10 - Receptor usage of PRCV-ISU1

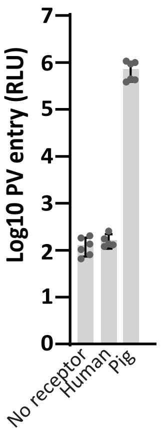

Fig S11 - Screening of human TMPRSS2 and human DPP4

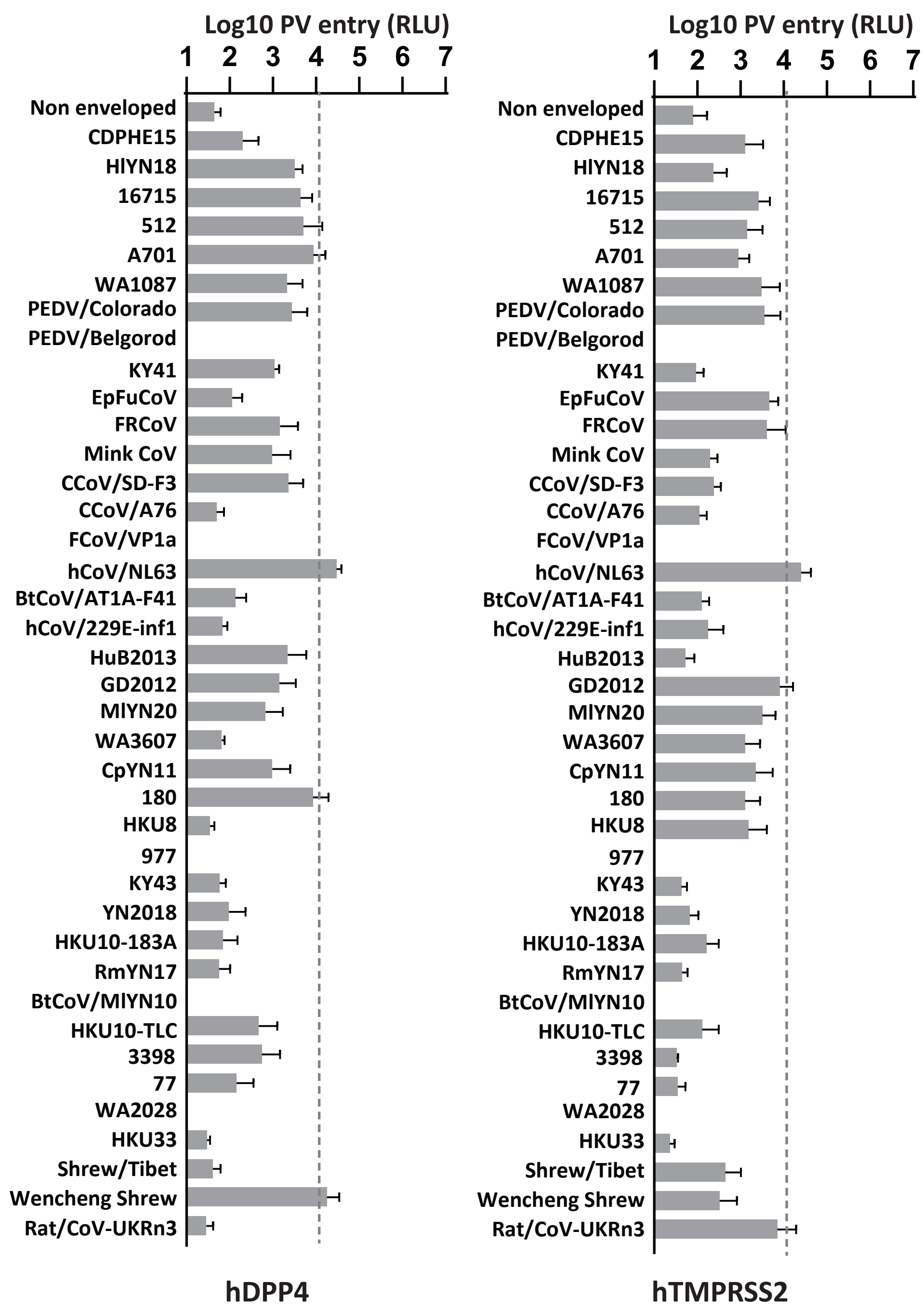

Fig S12 - Comparison of the binding interfaces of NL63 and SARS-CoV-2 RBDs on human ACE2

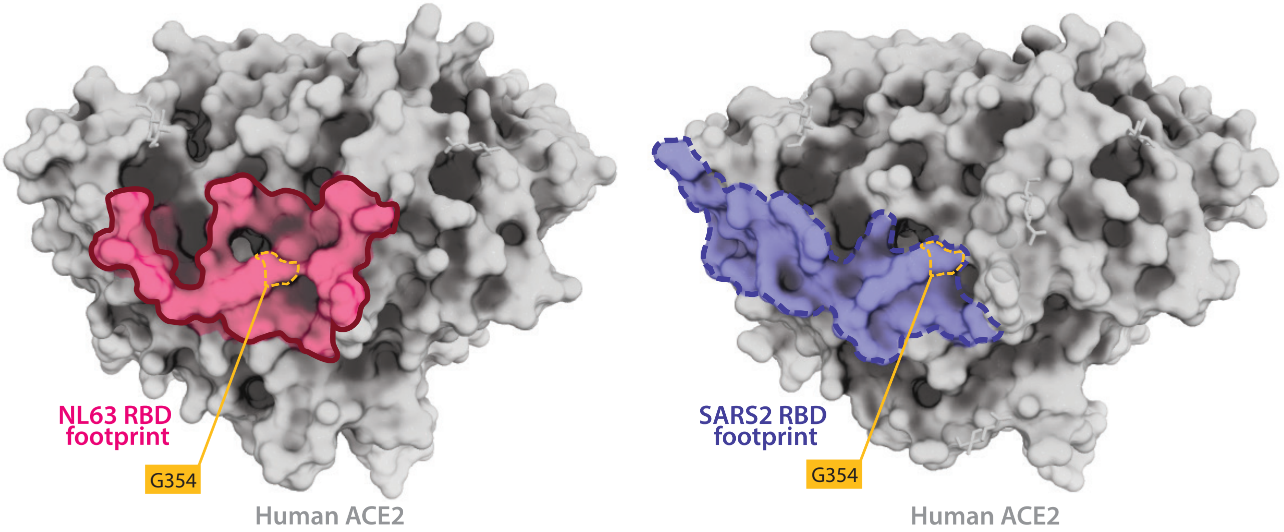

Fig S13 - APN screening of 229E-like mutants.

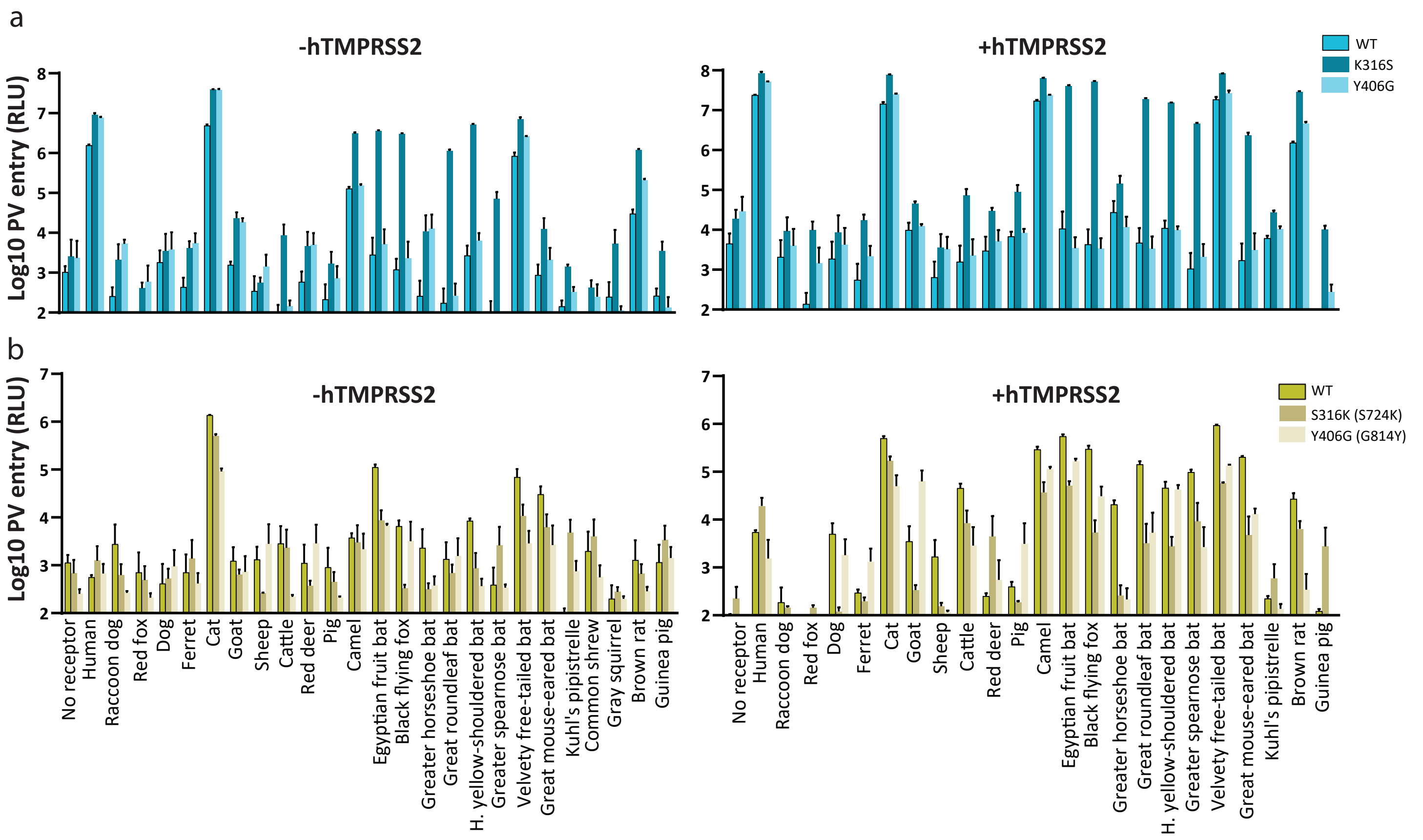

Fig S14 - APN screening of hCoV/229E, BtCoV/AT1A-F41 and BtCoV/WA1087 in presence of different human TMPRSS.

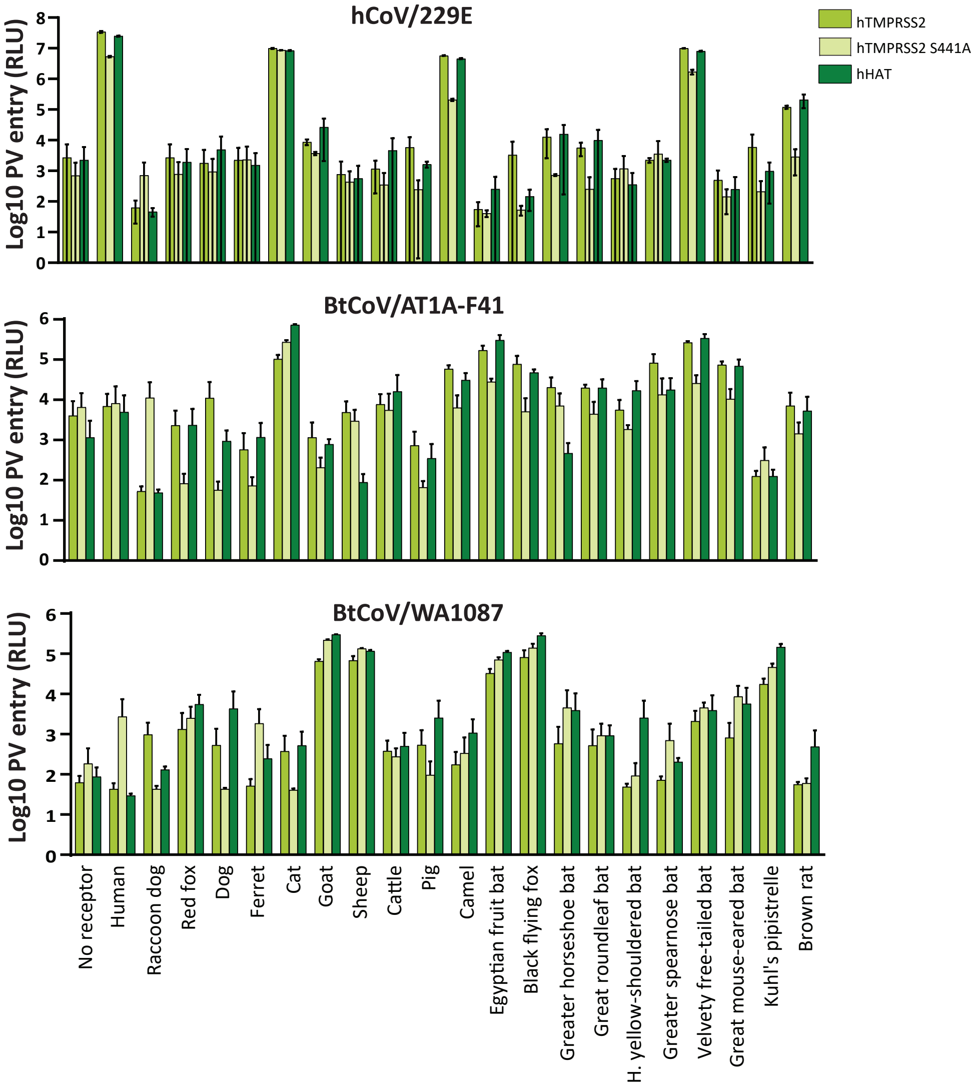

Table S1 - Genbank accession numbers of the alphaCoV S library

| ID | GenBank accession number |
|---|---|
| BtCoV/CDPHE15 | AGT21333 |
| BtCoV/HlYN18 | QWN56394 |
| BtCoV/16715 | AYR18455 |
| BtCoV/512 | YP-001351684 |
| BtCoV/A701 | ABG11965 |
| BtCoV/WA1087 | QGX41945 |
| PEDV/KDJ | AJD09595 |
| PEDV/Colorado | AGO58924.1 |
| PEDV/Belgorod | ASV51733 |
| BtCoV/KY41 | ADX59458 |
| EpFuCoV | UNE74462 |
| Ferret CoV | AKG92640 |
| Mink CoV | ADI80513 |
| CCoV/SD-F3 | ULF47968 |
| CCoV/A76 | AEQ61968 |
| FCoV/VP1a | UOS85847 |
| hCoV/NL63 | AIW52843 |
| BtCoV/AT1A-F41 | ALK28766 |
| hCoV/229E-ITA | QOP39313 |
| hCoV/229E-inf-1 | NP_073551.1 |
| BtMfCoV/HuB2013 | AIA62242 |
| BtMfCoV/GD2012 | AIA62212 |
| BtCoV/MlYN20 | QWN56401 |
| BtCoV/WA3607 | QGX41951 |
| BtCoV/CpYN11 | QWN56273 |
| BtCoV/180 | UIG55624 |
| BtCoV/HKU8 | ACA52171 |
| BtCoV/977 | ABO88151 |
| BtCoV/KY43 | ADX59451 |
| BtRsCoV/YN2018 | QDF43810 |
| BtCoV/HKU10-183A | YP-006908642 |
| BtCoV/RmYN17 | QWN56367 |
| BtCoV/MlYN10 | QWN56334 |
| BtCoV/HKU10-TLC | AFU92122 |
| BtCoV/3398 | YP-009755890 |
| BtCoV/77 | ULD45286 |
| BtCoV/WA2028 | QGX41957 |
| BtCoV/HKU33 | QCX35160 |
| Shrew-CoV/Tibet14 | ATP66784 |
| Wencheng Shrew-CoV | ASF90496 |
| Rat/CoV-UKRn3 | QBG64657 |
| BtCoV/MlYN15 | QWN56408 |
| CCoV/HuPn18 | QVL91811 |
| CCoV/Z19 | QWY12682 |

Table S2 - Genbank accession numbers of the APN library

| Common name | Scientific name | Genbank accession number |
|---|---|---|
| Human | *Homo sapiens* | NP_001141.2 |
| Raccoon dog | *Nyctereutes procyonoides* | XP_055182557.1 |
| Red fox | *Vulpes vulpes* | XP_025856768.1 |
| Dog | *Canis lupus familiaris* | NP_001139506.1 |
| Ferret | *Mustela putorius furo* | XP_012917463.1 |
| Cat | *Felis catus* | NP_001009252.2 |
| Goat | *Capra hircus* | XP_005695088.3 |
| Sheep | *Ovis aries* | XP_014957374.3 |
| Cattle | *Bos taurus* | NP_001068612.1 |
| European red deer | *Cervus elaphus* | OWK08899.1 |
| Pig | *Sus scrofa* | XP_005653580.1 |
| Bactrian camel | *Camelus ferus* | XP_006192640.2 |
| Egyptian fruit bat | *Rousettus aegypticus* | XP_016007055.1 |
| Black flying fox | *Pteropus alecto* | XP_006905412.1 |
| Greater horseshoe bat | *Rhinolophus ferrumequinum* | XP_032956109.1 |
| Great roundleaf bat | *Hipposideros armiger* | XP_019495552.1 |
| Honduran yellow-shouldered bat | *Sturnira hondurensis* | XP_036909514.1 |
| Greater spear-nosed bat | *Phyllostomus hastatus* | XP_045680750.1 |
| Velvety free-tailed bat | *Molossus molossus* | XP_03613394.1 |
| Greater mouse-eared bat | *Myotis myotis* | XP_036211683.1 |
| Kuhl's pipistrelle | *Pipistrellus kuhlii* | XP_36271433.1 |
| Common shrew | *Sorex araneus* | XP_004617569.1 |
| Gray squirrel | *Sciurus carolinensis* | MBZ3880029.1 |
| Brown rat | *Rattus norvegicus* | NP_112274.1 |
| Guinea pig | *Cavia porcellus* | XP_003475291.1 |

Table S3 - Genbank accession numbers of the ACE2 library

| Common name | Scientific name | Genbank accession number |
|---|---|---|
| Human | *Homo sapiens* | BAB40370.1 |
| Chimpanzee | *Pan troglodytes* | XP_016798468.1 |
| Rhesus macaque | *Macaca mulatta* | ACI04575.1 |
| Marmoset | *Callithrix jacchus* | XP_008987241.1 |
| Golden hamster | *Mesocricetus auratus* | XP_005074266.1 |
| Brown rat | *Rattus norvegicus* | NP_001012006.1 |
| House mouse | *Mus musculus* | NP_001123985.1 |
| Pig | *Sus scrofa* | NP_001116542.1 |
| Red deer | *Cervus elaphus* | XP_043752042.1 |
| Cattle | *Bos taurus* | NP_001019673.2 |
| Goat | *Capra hircus* | AHI85757.1 |
| Sheep | *Ovis aries* | XP_011961657.1 |
| Cat | *Felis catus* | AAX59005.1 |
| Ferret | *Mustela putorius furo* | BAE53380.1 |
| Mink | *Neogale vison* | XP_044091953.1 |
| Dog | *Canis lupus familiaris* | ACT66277.1 |
| Raccoon dog | *Nyctereutes procyonoides* | ABW16956.1 |
| Red fox | *Vulpes vulpes* | XP_025842513.1 |
| Little brown bat | *Myotis lucifugus* | XP_023609438.1 |
| Pallid bat | *Antrozous pallidus* | QJF77789.1 |
| Black-bearded tomb bat | *Taphozous melanopogon* | UJP38391.1 |
| Common vampire bat | *Desmodus rotundus* | XP_024425698.1 |
| Fruit bat | *Rousettus leschenaultii* | BAF50705.1 |
| Indian false vampire bat | *Megaderma lyra* | QKE49998.1 |
| Greater horseshoe bat | *Rhinolophus ferrumequinum* | BAH02663.1 |
| Halcyon horseshoe bat | *Rhinolophus alcyone* | ALJ94035.1 |
| Intermediate horseshoe bat | *Rhinolophus affinis* | QMQ39240.1 |
| Japanese horseshoe bat | *Rhinolphus cornutus* | BCG67443.1 |
| Shamel's horseshoe bat | *Phinolophus shameli* | UBB59645.1 |
| Pearson's Horseshoe bat | *Rhinolophus pearsonii* | QKE49996.1 |
| Chinese rufous horseshoe bat | *Rhinolophus sinicus* | QMQ39219.1 |
| Big-eared Horseshoe bat | *Rhinolophus macrotis* | ADN93471.1 |
| Indian flying fox bat | *Pteropus giganteus* | XP_039729365.1 |
| Least Horseshoe bat | *Rhinolophus pusillus* | ADN93477.1 |

Table S4 - ACE2 residues interacting with hCoV/NL63 RBD

| | ACE2 residues at 4.5Å from NL63 RBD (human ACE2 numbering) | | | | | | | | | |
|---|---|---|---|---|---|---|---|---|---|---|
| | 30 | 33-34 | 37 | 41 | 321-326 | 330 | 353-357 | 383 | 386-387 | 393 |
| Human | D | NH | E | Y | PNMTQG | N | KGDFR | M | AA | R |
| Chimpanzee | D | NH | E | Y | PNMTQG | N | KGDFR | M | AA | R |
| Macaque | D | NH | E | Y | PNMTQG | N | KGDFR | M | AA | R |
| Marmoset | D | NH | E | H | PNMTQG | N | KQDFR | M | AA | R |
| Syrian hamster | D | NQ | E | Y | PYMTQG | N | KGDFR | M | AT | R |
| Brown rat | N | NQ | E | Y | PQMTPG | N | HGDFR | M | AK | R |
| House mouse | N | NQ | E | Y | PHMTQG | N | HGDFR | M | AR | R |
| Pig | E | NL | E | Y | PNMTQG | N | KGDFR | M | AI | R |
| Red deer | E | NH | E | Y | PHMTQG | N | KGDFR | M | AA | R |
| Cattle | E | NH | E | Y | PYMTQG | N | KGDFR | M | AA | R |
| Goat | E | NH | E | Y | PYMTQG | N | KGDFR | M | AT | R |
| Sheep | E | NH | E | Y | PYMTQG | N | KGDFR | M | AT | R |
| Cat | E | NH | E | Y | PNMTQG | N | KGDFR | M | AV | R |
| Ferret | E | NY | E | Y | PNMTEG | N | KRDFR | M | AE | R |
| Mink | E | NY | E | Y | PNMTEG | N | KHDFR | M | AA | R |
| Dog | E | NY | E | Y | PNMTQE | N | KGDFR | M | AA | R |
| Raccoon dog | E | NY | E | Y | PNMTQG | N | RGDFR | M | AA | R |
| Red fox | E | NY | E | Y | PNMTQG | N | KGDFR | M | AA | R |
| Little brown bat | E | NS | E | H | PSMTPG | N | KGDFR | M | AT | R |
| Pallid bat | E | NS | E | H | PAMTPG | N | KGDFR | M | AS | R |
| Black-bearded tomb bat | E | NS | E | F | PNMTEG | N | QGDFR | M | ST | R |
| Common vampire bat | E | NT | E | Y | FSMTQG | N | NKDFR | M | AN | R |
| Fruit bat | E | NT | E | Y | PNMTET | K | KGDFR | M | AT | R |
| Indian flying fox | E | NT | E | Y | PNMTEK | K | KGDFR | M | AT | K |
| Indian false vampire bat | E | DL | E | Y | PNMTEG | N | KNDFR | M | AH | R |
| Greater horseshoe bat | D | NS | E | H | PNMTEG | N | KGDFR | M | AS | R |
| Halcyon horseshoe bat | D | NS | E | H | PHMTEG | N | KGDFR | M | AS | R |
| Intermediate horseshoe bat | D | NR | E | Y | PNMTEG | N | KGDFR | M | AT | R |
| Shamel's horseshoe bat | D | NP | E | H | PNMTEG | N | KNDFR | M | AT | R |
| Pearson's horseshoe bat | D | NR | E | H | PNMTEG | N | KDDFR | M | AS | R |
| Chinese rufous horseshoe bat | D | NS | E | Y | PNMTEG | N | KGDFR | M | AS | R |
| Big-eared horseshoe bat | D | NS | E | Y | PKMTEG | K | KGDFR | M | AS | R |
| Least horseshoe bat | N | NS | E | Y | PNMTEG | N | KGDFR | M | AS | R |
| Japanese horseshoe bat | N | NS | E | Y | PNMTEG | N | KGDFR | M | AS | R |

Table S5- APN residues interacting with hCoV/229E RBD

| | APN residues at 4.5Å from 229E RBD (human APN numbering) | | | | |
|---|---|---|---|---|---|
| | 286-292 | 303 | 309-310 | 315 | 318 |
| Human | EFDYVEK | W | IA | D | L |
| Raccoon dog | EFKNVQE | W | ID | N | L |
| Red fox | EFKNVQE | W | ID | N | L |
| Dog | EFKNVQE | W | MD | N | L |
| Ferret | EFKNLER | W | ID | N | L |
| Cat | EFSYVET | W | IN | D | L |
| Goat | EFTSVES | W | TA | L | L |
| Sheep | EFTSVES | W | TT | L | L |
| Cattle | EFTSVES | W | TA | L | L |
| European red deer | EFTSVES | W | TA | L | L |
| Pig | EFQSVNE | W | IA | M | L |
| Bactrian Camel | EFTCVEG | W | TA | I | L |
| Egyptian fruit bat | EFTCVQE | W | TK | L | L |
| Black flying fox | EFTCVEE | W | TR | L | L |
| Greater horseshoe bat | EFTSVEQ | W | TA | M | L |
| Great roundleaf bat | EFTAVEE | W | TK | I | L |
| Honduran yellow-shouldered ba | EFTSVER* | W | TQ | A | L |
| Greater spear-nose bat | EFTSVEQ* | W | TK | D | L |
| Velvety free-tailed bat | EFTHVEE* | W | TS | M | L |
| Greater mouse-eared bat | EFTYVNM* | W | TA | Q | L |
| Kuhl's pipistrelle | EFESVNS* | W | TE | Q | L |
| Common shrew | EFESVNS | W | IK | D | L |
| Eastern gray squirrel | EFKSVET | W | IE | E | L |
| Brown rat | EFKYVEA | W | ID | D | L |
| Guinea pig | EFHSVES | W | IA | E | L |

Table S6 - APN residues interacting with CCoV/HuPn18 RBD

| | APN residues at 4.5Å from HuPn18 RBD (canine APN numbering) | | | | | | | | |
|---|---|---|---|---|---|---|---|---|---|
| | 371-372 | 730 | 734 | 738-742 | 767 | 770-771 | 774-775 | 778 | 785-790 |
| Human | PL | F | R | NNWRE | E | EM | GL | Q | NNP-IHP |
| Raccoon dog | PQ | F | E | NNWTD | K | DL | TL | E | NNP-IYP |
| Red fox | PQ | F | E | KNWTD | K | DL | TL | E | NNP-IYP |
| Dog | PQ | F | E | QNWTD | K | DL | TL | E | NNP-IYP |
| Ferret | PL | F | E | QNWTK | K | EL | AL | E | NNS-IYP |
| Cat | RQ | F | E | KNWTD | E | KL | TL | Q | NNP-IHP |
| Goat | PQ | F | E | KGWTE | K | EL | TL | Q | ANP-INP |
| Sheep | PQ | F | E | KNWTE | K | EL | TL | Q | VNP-INP |
| Cattle | PQ | F | E | KNWTE | K | EL | TL | Q | VNP-IDP |
| European red deer | PQ | F | E | NNWTE | K | EL | TL | Q | VNP-IDP |
| Pig | PQ | F | E | KNWTE | Q | NL | TL | Q | NNP-IHP |
| Bactrian Camel | PL | F | E | KNWTE | K | EL | TL | S | NNP-IHP |
| Egyptian fruit bat | PL | F | K | KTWTQ | E | EL | NL | Q | NNP-IHP |
| Black flying fox | SL | F | K | KTWTQ | E | EL | SL | Q | NNP-IHP |
| Greater horseshoe bat | PQ | F | K | NKWTN | E | QL | EL | R | NNP-IHP |
| Great roundleaf bat | PE | F | Q | QNWTQ | E | QL | KL | D | NNP-IHP |
| Honduran yellow-shouldered bat | PL | F | R | RDWTQ | D | NL | QL | Q | SNP-IHP |
| Greater spear-nose bat | PQ | F | R | SDWTQ | G | TL | GL | Q | NNT-IHP |
| Velvety free-tailed bat | PA | F | E | KNWTQ | A | AL | RL | Q | NNP-IHP |
| Greater mouse-eared bat | PD | F | E | GNWTK | E | TL | DL | K | NNP-IHP |
| Kuhl's pipistrelle | PD | F | E | VNWTK | E | KL | DL | Q | INP-IHP |
| Common shrew | PG | Y | E | NNFTK | K | SL | EL | K | DKTIIHP |
| Eastern gray squirrel | PL | F | K | NNWKD | E | TL | SL | E | NNP-IHP |
| Brown rat | PQ | F | K | NNWLD | E | DL | GL | Q | NNP-IHP |
| Guinea pig | PE | Y | K | KEWSV | D | EM | NL | Q | NNP-IHP |

### Non pseudotyping with APN only (no more space in Fig. S5)

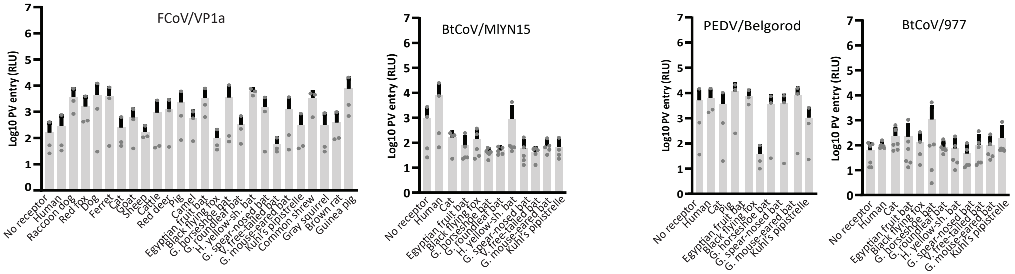
